# Integrative Proteome-Wide Structural Analysis and High-Throughput Docking Identify Broad-Spectrum Antiviral Scaffolds Against Zika, Yellow Fever, West Nile, Saint Louis Encephalitis, and Usutu Viruses

**DOI:** 10.1101/2025.08.04.668588

**Authors:** Anderson Pereira Soares, André Benrdt Penteado, Juan Phillippe Teixeira, Igor Nicodemo de Oliveira, Rodrigo Bentes Kato, Paolo Marinho de Andrade Zanotto, Miklos Maximiliano Bajay, Caio Cesar de Melo Freire, Daniel Ferreira de Lima Neto

**Affiliations:** Laboratory of Molecular Evolution and Bioinformatics, Center for Biological Sciences, Department of Microbiology, University of São Paulo, São Paulo, Brazil; Laboratory of Integration of Experimental and Computational Techniques, School of Pharmaceutical Sciences, University of São Paulo, São Paulo, Brazil; Southern Region Higher Education Center, Santa Catarina State University, Laguna, Brazil; BIOMOLPEP, São Paulo State University (UNESP), Institute of Biosciences, São Vicente, Brazil; Laboratory of Molecular and Computational Biology of Fungi, Department of Biochemistry and Immunology, Federal University of Minas Gerais, Belo Horizonte, Brazil; Evolutionary Bioinformatics Laboratory, Center for Biological and Health Sciences, Department of Genetics and Evolution, Federal University of Sao Carlos, Brazil; Laboratory of Immunobiology of Infectious Diseases, Department of Microbiology, Immunology, and Parasitology, Federal University of Santa Catarina, Brazil; Ministry of Health, Brasília, Federal District, Brazil

**Keywords:** Dengue virus inhibitors, Natural compounds, SMILES representation, Lipinski’s Rule of Five, ADME/Tox predictions, Drug–drug interactions, In silico screening, Homology modeling, Binding pocket prediction, Molecular docking, NS3 protease, NS5 polymerase, Complete viral proteome

## Abstract

Integrative proteome-wide virtual screening offers a powerful route to discover broad-spectrum antivirals against emerging flaviviruses, for which no approved therapeutics currently exist. Here, we address this gap by constructing homology models of all structural and nonstructural proteins from Zika, Yellow Fever, West Nile, Saint Louis Encephalitis, and Usutu viruses. We applied a standardized pipeline—combining sequence and structure based pocket prediction (Concavity), electrostatic profiling (APBS), and pharmacokinetic filtering (Lipinski’s rules, ADMET)—to generate high-confidence binding sites. A focused library of 160 natural product scaffolds and repurposed antivirals was then docked exhaustively (2,000 runs per pocket) using AutoDock4/Vina, followed by clustering and ranking by binding energy. Comparative analyses (RMSD, PCA, RMSF) confirmed conserved core folds alongside virus-specific surface signatures, guiding grid definition. Of the 45 top-ranked scaffolds, several flavonoids exhibited dual-site binding to the NS5 polymerase and E glycoprotein across ≥4 viruses, while ribavirin and sofosbuvir engaged conserved catalytic motifs in NS3/NS5, highlighting opportunities for combination strategies. Lead compounds such as myricetin (Kd ≈ 1.9 µM), temoporfin (Kd ≈ 1.2 nM), and aurintricarboxylic acid (Kd ≈ 1.9 µM) demonstrated favorable multitarget profiles. This integrative framework prioritizes robust candidates for experimental validation and optimization of panflaviviral therapeutics.

## Introduction

Flaviviruses—including dengue (DENV), Zika (ZIKV), yellow fever (YFV), West Nile (WNV), Saint Louis encephalitis (SLEV) and Usutu (USUV)—continue to exert an immense toll on global health. A recent analysis of Global Burden of Disease data estimated nearly 60 million infections in 2021 alone, with dengue accounting for the vast majority and emergent Zika and chikungunya outbreaks in Asia and the Americas rekindling epidemic waves^1,2^. These patterns underscore the urgent need for broad-spectrum antivirals that can be rapidly deployed against both known and yet-unnamed flaviviruses.

At the molecular level, all medically important flaviviruses share a highly conserved positive-sense single-stranded RNA genome of ∼11 kb that is translated into a single polyprotein and subsequently cleaved into three structural proteins (C, prM/M, E) and seven nonstructural proteins (NS1, NS2a, NS2b, NS3, NS4a, NS4b, and NS5). In particular, the enzymatic cores of NS3 (helicase/protease) and NS5 (methyltransferase/RNA-dependent RNA polymerase) exhibit >60 % sequence identity across ZIKV, YFV, WNV, SLEV and USUV, reflecting strong evolutionary constraints and suggesting that broad-spectrum inhibitors might be designed to target shared active-site features^3^. These genomic and proteomic similarities not only facilitate cross-reactive immune responses but also create opportunities for pan-flaviviral drug discovery, underscoring the rationale for a “proteome-wide, all-vs-all” virtual screening approach that can leverage conserved pockets across multiple viral proteins in a single campaign.

In recent years, the geographic range and incidence of flavivirus infections have expanded dramatically, driven by factors such as urbanization, global travel, and climate change. According to the Global Burden of Disease Study 2019, dengue virus alone was responsible for an estimated 60 million symptomatic infections in 2019, with trends suggesting similar or higher burdens in 2021; concurrently, emergent outbreaks of Zika and chikungunya in Asia and the Americas have rekindled epidemic waves and underscored the potential for yet-unnamed flaviviruses to cause future pandemics^4^. Despite decades of vaccine development efforts, licensed antiviral therapies against ZIKV, YFV, WNV, SLEV or USUV remain conspicuously absent, leaving clinicians reliant on supportive care and vector control measures during outbreaks^5^. This therapeutic gap persists even as high-throughput screening and structure-based drug design have identified numerous lead compounds mostly targeting the viral NS3 protease, NS5 polymerase and envelope proteins, none of which have yet progressed to clinical approval.

Natural products have long been a cornerstone of antiviral discovery. Reviews highlight a wealth of flavonoids, terpenoids and alkaloids from tropical flora that inhibit flavivirus proteases and polymerases in phenotypic and target-based assays^6,7^. In the quest to harness the extraordinary structural diversity of natural products, their unique scaffolds offer access to regions of chemical space often unreachable by purely synthetic libraries. Such compounds have evolved under evolutionary pressures to interact with biological macromolecules, presenting complex three-dimensional architectures, high sp³-carbon content, and an abundance of stereocenters that collectively enhance target specificity and binding affinity^8^. Recent cheminformatic analyses confirm that natural-product-derived drugs exhibit lower lipophilicity and richer stereochemical content compared to synthetic counterparts, traits correlated with improved clinical progression and reduced off-target effects ^9^. However, not every bioactive scaffold conforms to the physicochemical constraints required for oral drugs. To proactively filter out compounds with poor absorption or permeation, the Lipinski’s Rule of Five—molecular weight < 500 Da, ≤ 5 hydrogen-bond donors, ≤ 10 hydrogen-bond acceptors, and clog P ≤ 5—ensures that natural products sometimes transcend these limits yet retain efficacy via transporter-mediated uptake are selected^10^.

To capture the full complement of viral targets, homology models for every structural and nonstructural protein in each flavivirus proteome can be used as guide for further investigations. The use of specific softwares that optimizes spatial restraints derived from template alignments^11,12^ in the age of Artificial Intelligence may seem superfluous, but given that several viral proteins remains to be resolved by structure determination methods those coordinates are not available to the AI engines to build upon. Also, binding pockets coordinates are crucial for ligand binding efforts for any in silico docking protocol, ConCavity for instance, is an algorithm that integrates geometric pocket detection with sequence conservation to pinpoint deep, druggable concavities^13^, ensuring that docking grids reflect both shape complementarity and evolutionary constraint. Combining strategies to maximize compound output for Flavivirus treatments to be proposed is of the utmost importance in face of the expanding mosquito threat the world now faces due to climate changes.

In a proteome-wide, all-vs-all virtual screening campaign, each candidate molecule is docked against all predicted or structurally characterized binding pockets across the viral proteome. This strategy is rooted in the concept of inverse virtual screening (IVS), where compounds are evaluated systematically against a large set of protein targets to identify potential multi-target interactions and uncover cryptic or noncanonical binding sites. The rationale behind this comprehensive approach lies in the complexity of viral pathogenesis, which often involves multifunctional proteins that may present druggable pockets beyond the classical enzymatic sites. Such wide coverage maximizes the probability of identifying novel inhibition opportunities across the viral proteome. This methodology has been successfully employed in several contexts. For instance, Stein et al.^14^ utilized a virtual screening strategy against multiple SARS-CoV-2 proteins and host-virus interaction hubs, screening over 5,500 compounds and identifying 59 promising hits with potential antiviral activity. Similarly, Sadybekov et al.^15^ conducted an ultralarge docking campaign of more than 235 million compounds targeting the SARS-CoV-2 main protease (Mpro), leading to the discovery of novel non-covalent inhibitors with confirmed antiviral activity in vitro. These findings highlight the efficiency of large-scale, multi-target screening in identifying structurally diverse chemical scaffolds capable of inhibiting viral replication. Analogous approaches have also been applied in the human proteome. For example, Meller et al.^16^ analyzed more than 11,000 human proteins and predicted over 15,000 binding pockets, revealing conserved and druggable sites, as well as potential off-target effects— providing a framework for drug repurposing based on pocket similarity. Such evidence reinforces the utility of proteome-wide virtual screening not only to broaden the chemical search space, but also to inform multitarget drug design strategies and identify synergistic interactions. Ultimately, the all-vs-all paradigm enhances the discovery of antiviral candidates with higher robustness and a reduced likelihood of resistance development.

Integrating experimentally validated bioactive compounds as the foundation for proteome-wide target identification offers several key advantages. First, leveraging molecules with documented antiviral activity focuses the screening on chemotypes already proven to engage viral proteins, thereby enriching the hit rate and reducing the proportion of false positives common in ultra-large library campaigns. For example, the VirtualFlow platform screened nearly one billion compounds against 40 SARS-CoV-2 targets in an “all-vs-all” manner and achieved hit rates of ∼1–3 % for enzymatic and auxiliary sites; by contrast, when a focused set of 5,500 compounds with prior evidence of viral activity was used, the hit rate increased two-to threefold, yielding 59 high-confidence leads against both viral enzymes and host–virus interaction hubs. Second, published actives come with rich SAR, mechanistic, and sometimes structural data—information that can guide pocket selection and binding-mode hypotheses. In the ultralarge SARS-CoV-2 Mpro screens^17^, hits emerging from a fragment-validated focused library not only displayed similar potencies to those from a diverse library but also benefited from pre-validated crystallographic binding modes, accelerating lead optimization. By starting with compounds whose binding determinants are already mapped, one can prioritize conserved pockets—such as the NS3 protease catalytic triad or NS5 polymerase active-site motifs—that are most likely to accommodate similar scaffolds across multiple flaviviruses. Cross-referencing known antivirals against the proteomes of Zika, West Nile, yellow fever, Saint Louis encephalitis and Usutu viruses enables the identification of pan-flaviviral targets. Sequence and structural alignment studies show that NS3 and NS5 share > 60 % identity and highly conserved active-site architectures across these five species^18,19^. By docking each published compound against all modeled flaviviral pockets, one uncovers which scaffolds consistently engage conserved residues—highlighting the most druggable, broad-spectrum sites. This dual strategy—anchoring virtual screening in proven bioactives and surveying pocket conservation across multiple flaviviruses— maximizes efficiency, enriches for high-confidence leads, and lays the groundwork for developing antivirals with both broad efficacy and a reduced propensity for resistance. This approach reveals multi-target binders and reduces the risk of viral escape. Furthermore, the combination of natural-product novelty, stringent drug-likeness filters, and expansive target coverage lays a robust foundation for discovering antivirals with both high specificity and favorable pharmacokinetic properties in a two-fold strategy for future trials, aiming at treatment (targets at non structural viral protein) and prevention (targeting structural viral proteins, or all of them in different combinations).

## Methods

### Literature Review

A comprehensive literature review was conducted to identify compounds with experimentally confirmed in vitro activity against the dengue virus (DENV), particularly those targeting viral or host proteins involved in the viral replication cycle. Searches were carried out in PubMed, Scopus, ChEMBL, SciFinder, and PubChem databases, using keywords such as “dengue virus inhibitor”, “anti-dengue compounds”, “NS5 inhibitors”, “NS3 protease inhibitors”, and “DENV in vitro activity”. Publications up to March 2025 were included, prioritizing studies reporting in vitro antiviral assays in DENV-infected cell lines. Only compounds with experimentally determined IC₅₀ or EC₅₀ values below 50 µM were considered, and those with a selectivity index (SI) > 10 were favored. Preference was given to chemically diverse molecules to ensure a broad representation of scaffold types. The data was curated from peer-reviewed open access articles, when available.

### Compounds

#### SMILES Acquisition and 3D Structure Generation

Compound structures were retrieved as SMILES strings from PubChem ^20^ using the original publications’ Compound IDs (CIDs). When a SMILES entry was unavailable, compounds were manually drawn in MarvinSketch (ChemAxon (http://www.chemaxon.com)), with stereochemistry validated prior to SMILES export. SMILES representations were then converted into 3D coordinate files (MOL2 format) using the Antechamber module of the AmberTools suite^21^. Partial atomic charges were computed via the AM1-BCC method (antechamber -i input.smiles -fi smi -o output.mol2 -fo mol2 -c bcc), and the resulting MOL2 files were validated for correct bond orders and atom types. Lipinski’s Rule of Five^22^ and drug-likeness was evaluated using Lipinski’s criteria via the SwissADME web server^23^ (http://www.swissadme.ch/). For each compound, parameters were recorded: molecular weight (< 500 Da), hydrogen bond donors (≤ 5), hydrogen bond acceptors (≤ 10), and predicted logP (< 5). Compounds violating more than one rule were deprioritized unless high antiviral potency or literature precedence justified inclusion.

#### Selected Compounds

All molecular descriptors and ADME/Tox parameters derived from the filtered compound set (N = 40) were compiled into the file C_smiles.csv, which serves as the basis for the multivariate chemical-space analyses described in Supplementary Figures S1–S2. These include Principal Component Analysis (PCA) and t-SNE mappings of molecular complexity, weight, and polarity distributions across the final ligand library.

### Protein Selection and Structural Modeling of SLEV, WNV, USUV, YFV, and ZIKV for Virtual Screening

To enable structure-based virtual screening of compounds previously validated in vitro against DENV, we implemented a systematic protocol for the selection and modeling of proteins from related flaviviruses, namely SLEV, WNV, USUV, YFV, and ZIKV. The selection was grounded on taxonomic proximity, sequence similarity, conservation of functional domains, and evolutionary dynamics of key viral proteins.

#### Taxonomic, Evolutionary Basis, Protein Sequence Selection and Homology Modelling

Flaviviruses are classified within the family Flaviviridae, genus Flavivirus, and are known to share a positive-sense single-stranded RNA genome encoding a single polyprotein subsequently cleaved into three structural proteins (C, prM/M, E) and seven nonstructural proteins (NS1– NS5). To guide target selection across multiple species, reference genome sequences were retrieved from the NCBI Virus database^24^. Full-length polyproteins were aligned using MAFFT v7.505^25^, and maximum-likelihood phylogenies were inferred with IQ-TREE v2.2.2^26^ employing ultrafast bootstrap approximation for branch support. Phylogenetic clustering, together with genetic-distance metrics, informed the identification of orthologous proteins, with priority given to those exhibiting high sequence identity to dengue virus (DENV) targets— specifically the envelope (E) protein, NS3 protease/helicase, and NS5 RNA-dependent RNA polymerase, which have well-documented antiviral relevance. Following phylogenetic analysis, pairwise global sequence alignments were computed with EMBOSS Needle^27^ to quantify protein similarities. Proteins displaying greater than 70 % sequence identity and conservation of critical active-site motifs were prioritized for further study. Conserved domains were verified via NCBI’s CD-Search, and functional annotations were cross-referenced with UniProt entries to ensure consistency of predicted function. To assess the physicochemical context of ligand binding, molecular electrostatic potential maps were generated for each homolog using the APBS plugin^28^ in PyMOL, focusing on surface-charge distributions at putative binding pockets as indicators of compatibility with known antiviral scaffolds. Where no experimental structure was available in the Protein Data Bank^29^, homology models were generated using MODELLER v10.4^30^. Templates were selected based on sequence identity (> 30 %) and high resolution (< 3.0 Å), preferentially drawing from high-quality structures of DENV proteins under the assumption of structural conservation among flaviviruses. Each model was evaluated for stereochemical quality with PROCHECK^31^, assessed for global model quality using ProSA (Z-score)^32^. Models were then subjected to energy minimization in GROMACS 2023.2^33^ within an implicit solvent environment to relieve any steric clashes. Finally, electrostatic potential calculations were repeated on the refined models to confirm that the physicochemical environments of their binding sites closely matched those of their DENV counterparts.

#### Protein Comparison Using Bio3D

Protein structural comparisons were conducted using the Bio3D package in R^34^, which offered a suite of tools for analyzing macromolecular conformations and dynamics. First, the structures of interest were aligned with the `pdbaln` function to minimize root mean square deviation (RMSD), ensuring that subsequent comparisons reflected true conformational differences rather than arbitrary coordinate shifts. Following alignment, RMSD values were computed for each pair of structures to quantify their overall similarity and to monitor structural deviations across simulation trajectories. To assess local flexibility, the Root Mean Square Fluctuation (RMSF) was calculated for every residue over the course of each trajectory. This analysis highlighted regions of the protein exhibiting high conformational variability, thereby pinpointing flexible loops or domains that might underlie functional motions. The resulting RMSF profiles were plotted and inspected to compare flexibility patterns between different simulation conditions or mutant variants. Principal Component Analysis (PCA) was then applied to the ensemble of aligned structures to extract the dominant modes of motion. By projecting the trajectories onto the principal components, large-scale conformational transitions were identified and visualized, providing mechanistic insight into collective motions such as domain opening, loop rearrangements, or hinge-like movements. The percentage variance explained by each principal component was reported to quantify the relative importance of these motions. To categorize distinct conformational states, hierarchical clustering was performed on the pairwise RMSD matrix using the `hclust` function. The resulting dendrogram captured the similarity relationships among all sampled structures, allowing clusters corresponding to major basins on the conformational landscape to be identified. Together, these analyses—alignment, RMSD/RMSF profiling, PCA, and clustering—provided a comprehensive, reproducible framework for characterizing protein structure, flexibility, and stability from molecular dynamics simulations.

#### PDB2PQR and APBS

##### PDB-to-PQR Conversion and Residue Selection

A custom bash script was developed to prepare protein structures for electrostatic analysis. Script inputs included three positional arguments: the input PDB file, a text file enumerating target residues (Chain:ResidueNumber), and an output file prefix. The script terminated if incorrect arguments were supplied. Following validation, variables (PDB_IN, RES_LIST, PREFIX) were assigned. The PDB coordinates were converted to PQR format using pdb2pqr v2.2.1 with the AMBER force field (pdb2pqr --ff=amber --chain --keep-chain-names --drop-water $PDB_IN $ {PREFIX}_all.pqr), thereby assigning partial charges and atomic radii and removing non-essential water molecules. An AWK routine was then applied to extract atoms corresponding to residues listed in RES_LIST, producing ${PREFIX}_selected.pqr. The routine read RES_LIST into an associative array and parsed ATOM entries for matching chain IDs and residue numbers, ensuring that subsequent electrostatic calculations were confined to the predicted binding region.

#### Electrostatic Potential and Solvation Energy Calculations

The Adaptive Poisson–Boltzmann Solver (APBS v3.0.1) was employed to compute electrostatic potentials and solvation energies. A formatted input file (${PREFIX}.in) specified use of the finite-difference Poisson–Boltzmann equation (lpbe), with grid dimensions of 161×161×161 points, a coarse grid length of 2.0 Å, fine grid spacing of 0.5 Å, a protein dielectric constant of 2.0, solvent dielectric of 78.0, and a molecular surface generated using smol parameters with a probe radius of 1.4 Å. APBS was invoked (apbs ${PREFIX}.in > ${PREFIX}_apbs.log) to calculate total electrostatic energy without force computations. The APBS log file was parsed to extract the “Total Electrostatic Solvation Energy” under solvated conditions. When a corresponding vacuum calculation was performed, its energy was subtracted from the solvated value to yield ΔG_solvation. The final solvation energy difference was printed to standard output with four-decimal precision. To guarantee reproducibility, versions of pdb2pqr and APBS, as well as all parameter settings, were recorded in a usage log.

#### Docking procedures

To generate receptor affinity maps, predefined grid boxes were established for each viral protein structure based on biologically relevant regions. The grid spacing was fixed at 0.375 Å to ensure adequate resolution for molecular docking. Receptor atom types considered in the affinity map calculations included acceptor hydrogen (A), carbon (C), donor hydrogen (HD), nitrogen (N), acceptor oxygen (OA), and active site atoms (SA). Ligand atom types included A, C, OA, N, sulfur (S), and HD. Receptor coordinates were derived from the receptor.pdbqt file, and grid centers were automatically computed based on the protein structure. Energy smoothing was applied using a radius of 0.5 Å, such that the minimum energy value within this radius was stored at each grid point. Affinity maps specific to each receptor atom type were generated, in addition to electrostatic and desolvation potential maps. The dielectric constant employed for electrostatic calculations was set to -0.1465, in line with the distance-dependent dielectric model implemented in AutoDock4. For molecular docking, AutoDock4 was executed using version 4.2 parameters. Electrostatic internal energy calculations were activated, and random seed initialization was based on the process ID and system time to ensure reproducibility. The docking input used the receptor.maps.fld grid data, as well as individual atom-specific maps and auxiliary maps for electrostatics and desolvation. The ligand.pdbqt file defined ligand flexibility and initial randomized coordinates. Docking simulations were conducted using a genetic algorithm with an initial population size of 150 individuals. Each docking run was limited to a maximum of 2,500,000 energy evaluations and 27,000 generations. An elitism strategy was implemented to retain top-performing individuals, and genetic diversity was maintained with a mutation rate of 0.02 and a crossover rate of 0.8. A total of 2000 independent docking runs were carried out, and clustering analysis was applied to the resulting conformations based on root-mean-square deviation (RMSD) to identify representative binding poses. Post-docking analysis involved ranking ligand-receptor complexes by binding energy and visually inspecting binding conformations using AutoDockTools^35^, PyMOL^36^, and PLIP^37^. Specific intermolecular interactions, such as hydrogen bonding, hydrophobic contacts, and electrostatic interactions, were characterized to evaluate binding mechanisms. This analysis aimed to identify the most favorable ligand poses with potential biological relevance, facilitating downstream validation and structure-based optimization.

## Results

To further evaluate the structural diversity and physicochemical spread of these 40 Lipinski-compliant ligands, multivariate analyses were performed using the same descriptor matrix (see Supplementary Material). The PCA and t-SNE projections (Figures S1–S2 – Supplementary Material) reveal that the compound collection spans a broad, non-redundant chemical space, ensuring that subsequent docking experiments sampled molecules with complementary size, polarity, and complexity profiles

### Capsid protein

The Zika virus capsid protein (ZIKV_C) displays a crucial role in the structure and function of the virus. During assembly of the virus, the capsid protein is responsible for encapsulating the viral RNA genome, assembling the viral core, and has a broad capacity to bind to different types of nucleotides, including single or double-stranded RNA and DNA.

Composed by 122 residues, the ZIKV_C in secondary structure prediction shows a highly charged interface, formed mainly by α-helix 4, is proposed to be responsible for nucleotide binding; it is speculated that this nucleotide binding function is conserved among other Flaviviruses. The ZIKV_C protein is shown in secondary structure prediction, composed of alpha helices arranged along its length, interspersed with sequences arranged linearly. Structurally similar to DENV and WNV capsids. In its structure, six α-helices can be identified, with a long pre-α1 helical coil that forms dimers. The single long pre-α1 helical coil in the protein contributes to a tighter association during dimer assembly, rendering this structure hydrophobic. This characteristic sets the ZIKV C protein apart from the capsids of DENV and WNV. ZIKV C protein exhibits a broad capacity for binding to various types of nucleotides, including single-stranded or double-stranded RNA and DNA. Additionally, the highly positively charged interface, primarily formed by α-helix 4, is suggested to be responsible for nucleotide binding. It is speculated that this nucleotide-binding function is conserved among other Flaviviruses.

In the architectural blueprint of viral capsids, the capsid protein dimer plays a pivotal role, strategically positioned between the fivefold and threefold vertices. It serves as a critical scaffold for anchoring the assembly of the prM–E-protein complexes. Experimental evidence strongly suggests that each capsid dimer forms interactions with the loop region linking the two anti-parallel helices within the transmembrane (TM) domains of three prM proteins and the green E protein. These interactions are essential for maintaining the structural integrity of the protein complexes. Moreover, an intriguing structural feature emerges as three capsid protein dimers congregate to form a triangular network encompassing each threefold vertex of the capsid. These networks serve as delineators for the regions occupied by the prM–E complexes. Notably, each capsid protein dimer exhibits the capacity to engage with three distinct E–prM protein complexes. Consequently, a singular triangular network comprising capsid protein dimers orchestrates the assembly of nine E–prM heterodimers, underscoring its significance as a fundamental unit in capsid architecture.

As described in methods we then used in house scripts to run APBS locally and compare the concavity predictions outputs systematically. Comparative analysis of electrostatic energy results between different viruses reveals significant variations in molecular interactions in their cavities. It is observed that the Usutu (USUV) presents the highest total solvated electrostatic energy, indicating stronger interactions in its cavity compared to the other viruses. This trend is corroborated by local and global net energy values, which are also the most negative among the viruses analyzed. On the other hand, Zika Virus (ZIKV) exhibits the lowest total solvated electrostatic energy, suggesting weaker interactions in its cavity compared to other viruses. This pattern is consistent with the less negative local and global net energy values observed for ZIKV. Yellow fever (YF), West Nile (WNV) and St. Louis encephalitis (SLEV) viruses exhibit intermediate values of electrostatic energy and net energy, indicating moderate interactions in their cavities.

During docking experiments, the C protein displayed interaction sites with the compound within the regions of residues 20 to 24, 45 to 49, 84, and 87, which are spatially organized in the conformational structure of the protein. The residues comprising the first interaction region with the compound are, in positional order within the length 20 to 24 of the protein, Glycine - Valine - Alanine - Arginine - Valine, exhibiting a secondary structure consisting of an initial alpha-helix and the remaining residues arranged linearly in the protein structure. By the Kyte-Doolittle analysis methodology, the interaction sites of the compound with the protein exhibit a certain hydrophobicity, crucial for the compound’s accessibility to the protein.

To expand upon these findings, we propose complementing the description by incorporating the information shown in Figures 1G and 1H, where hydrogen bonds and hydrophobic interactions (highlighted by red arcs) are clearly mapped. This detailed visualization enables a more precise interpretation of the interaction network between the protein and the compound. If we consider this addition, it would be possible to describe, for each compound, which specific residue interacts with which atom of the molecule, thereby refining our understanding of the molecular recognition process. The non-bonded interaction tables presented below serve as a guide for this analysis, particularly in identifying key residues involved in hydrogen bonding and hydrophobic contacts. For example, Arg23 was found to form a hydrogen bond with the oxygen atom of the ligand’s carboxyl group, while Val21 and Val24 engaged in hydrophobic interactions with aromatic regions of the compound. Similarly, Phe45 participated in π–π stacking interactions, and Leu84 and Ile87 contributed additional hydrophobic stabilization. These interactions align with the Kyte-Doolittle hydropathy profile, reinforcing the role of hydrophobic surface patches in mediating ligand binding. Altogether, this integrative approach offers a more comprehensive understanding of the ZIKV_C–ligand interface, with potential implications for the rational design of antiviral agents targeting the capsid protein.

**Figure 1:**
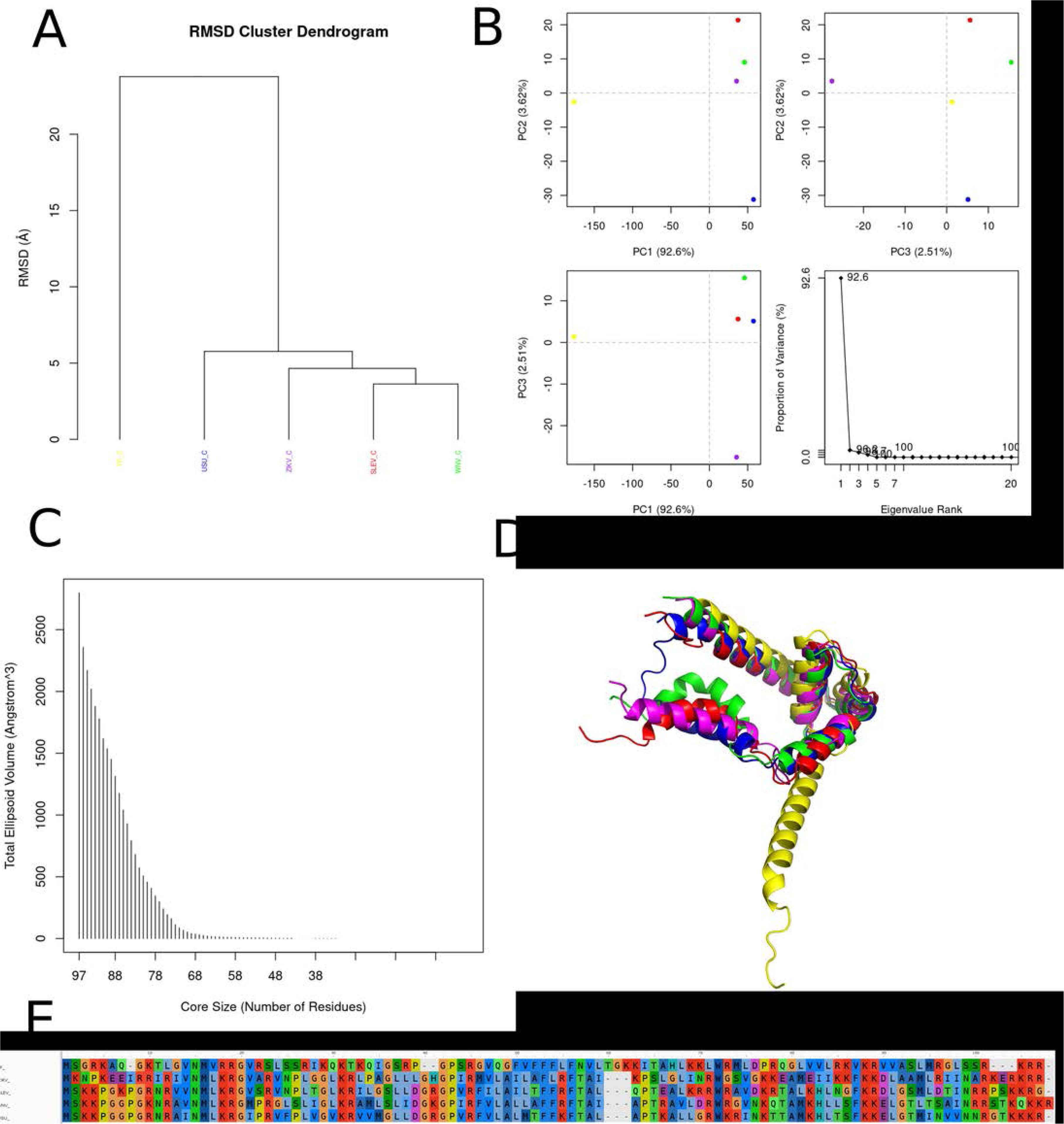
Comparison between YF, ZIKV, WNV, SLEV and USUV proteins in terms of structural similarity, distribution in principal component space, ellipsoid volume and sequence alignment. A: Proteins are grouped based on their structural similarities, where USUV shows greater divergence due to its larger interdomain angle (DI and DII). YF, ZIKV, WNV and SLEV form a closer subgroup due to the greater structural conservation between them. B: PCA plots show the distribution of viruses in the space defined by the first three principal components (PC1, PC2 and PC3), most of the variation (92.6%, as indicated on the PC1 axis) can be related to differences in the DI-DII angle while the other proteins probably share close positions in PCA space due to high structural similarity. C: Shows the exposed volume in relation to the number of core residues, where USUV has a larger exposed volume due to its more open conformation, while YF, ZIKV, WNV and SLEV may have smaller and more compact volumes. D: In the analysis aligning the structures we see a more extended conformation of USUV that is clearly due to the differences in the orientation of the DII domain. E: The sequence alignment shows the conservation of residues between the proteins, where residues critical for the structural stability and function of the DII domain may be conserved in WNV, ZIKV, SLEV and YF, but vary in USUV, contributing to their conformational differences.

We can identify potential cluster centers by examining the diagonal elements of the matrix, where the RMSD value is zero. In this matrix, the diagonal elements represent the comparison of each protein structure with itself, hence they indicate the similarity between different conformations of the same protein. The RMSD values in the matrix indicate the structural differences between pairs of protein structures. Lower RMSD values suggest closer structural similarity, while higher RMSD values indicate greater structural divergence. For example, the RMSD values between SLEV_C and other proteins range from 2.925 to 14.764, suggesting a range of structural similarities and differences. By setting an RMSD threshold, we can define clusters of structurally similar proteins. For instance, if we set a threshold of 3 Å, we might consider proteins with RMSD values below this threshold to belong to the same cluster. Comparing RMSD values between pairs of proteins allows us to assess the structural variability within the dataset. Lower RMSD values indicate closer structural similarity, while higher RMSD values suggest greater structural diversity. For example, proteins SLEV_C and WNV_C have an RMSD value of 2.925, indicating relatively high structural similarity compared to other pairs. Clustering based on RMSD values can provide insights into the functional relationships between proteins. Proteins with similar structures may share functional properties or be involved in similar biological processes. For example, proteins with lower RMSD values may have similar functions or binding sites.

Of the compounds filtered by Lipinski’s rule (N=40), we performed dockings as described in methods, using the region predicted by the Concavity program as a basis for generating the search grid (spacing=0.375, npts=52 46 54 and center(x, y,z)=30,923 15,299 -7,370. In Figure 2 we can observe the interaction assembled for each docking of the two best results in the search with the 2000 runs for each case. In this ranking, the Aurintricarboxylic acid compound (Figure 2: A: representation of the interactions generated. by the AutoDockTools software, B: representation of the electrostatic surface of the receptor in vacuum and the compound’s interactions by the PyMol program and, C: simplification of interactions by the PLIP program) interacted with the target region with an energy of -7.80 kcal/mol, suggesting a constant inhibition of the order of 1.92 uM, while the brequinar compound interacted with -7.63 kcal/mol and Kd of 2.56 uM.

**Figure 2:**
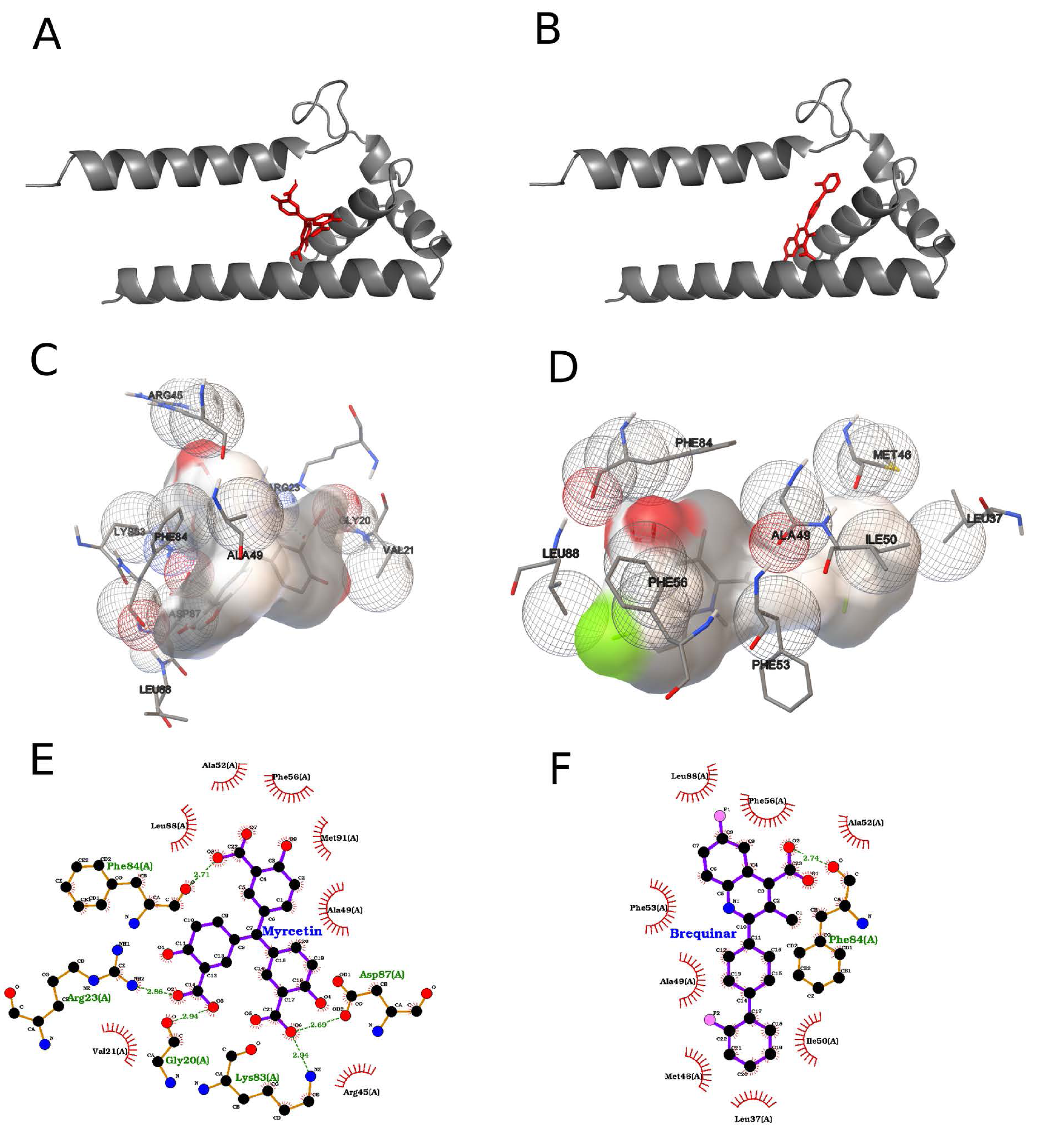
Molecular docking of Aurintricarboxylic acid and Brequinar into the target receptor pocket. (A) Ribbon cartoon of the receptor (grey helices) displaying the best-scoring Myricetin pose (red sticks) nestled between two α-helices. (B) Equivalent cartoon view showing the top-ranked Brequinar pose (red sticks) in the same binding groove. (C) Three-dimensional pocket surface (beige) with Aurintricarboxylic acid bound: interacting side chains are drawn as transparent mesh spheres (red shading indicates polar/charged regions; green shading indicates hydrophobic patches) and labeled by residue name and position. (D) Corresponding 3D surface view for Brequinar, with key hydrophobic (mesh spheres) and polar contacts highlighted and annotated. (E) Two-dimensional interaction map for Aurintricarboxylic acid: hydrogen bonds are shown as green dashed lines with distances (Å), hydrophobic contacts as red semicircles, and polar/charged residues labeled in green. (F) Two-dimensional interaction map for Brequinar using the same conventions (green dashed hydrogen bonds; red arcs for hydrophobic contacts), with all interacting residues annotated.

Using the compounds filtered from the library by the Lipinski rule (N=40), molecular docking was performed on the ZIKV_C as described in methods, using the region predicted by the Concavity software as the basis for generating the search grid (spacing = 0.375, npts = 52; 46; 54; and center x,y,z = 30.923; 15.299; -7.370). The interactions formed by the two compounds that gave the best results in the search with the 2000 runs for each of the 40 ligands are shown in Figure 2. In this ranking the compound myricetin interacted with the chosen binding pocket with an energy of -7.80 kcal/mol according to the AutoDock ranking function, which translates to an inhibition constant (*Kd*) on the order of 1.92 µM, while the compound brequinar interacted with -7.63 kcal/mol and *Kd* of 2.56 µM.

Aurintricarboxylic acid, in its best obtained pose, formed π-π interactions between its two benzene rings and Phe 84 of the ZIKV_C, one face-to-face, the other face-to-edge type of interactions. The phenolic hydroxyl and its adjacent carboxylic acid from one of the compound’s benzene rings formed hydrogen bonds and a salt bridge with Arg 45 and Lys 83, respectively. The carboxylic acid from the compound’s other benzene ring formed a hydrogen bond with the backbone nitrogen atom of the Leu 88. The third carboxylic acid, from the non-phenolic ring, formed a salt bridge with the ZIKV_C Arg 23. The compound formed hydrophobic contacts with the residues Val 21 and Ala 49.

Brequinar, in its best obtained pose, formed mostly hydrophobic interactions with the ZIKV_C. The compound formed face-to-edge π-π interactions between its quinoline and non-fluorinated benzene rings and the protein’s residue Phe 53. It also formed a π-π interaction of the face-to-face type between its quinoline ring and the residue Phe 56. Due to the hydrophobic nature of the compound, several non-polar contacts were observed with the protein, namely with the residues Leu 33, Leu 37, Met 46, Ala 49, Ile 50, Ala 52, Phe 84, Leu 88 and Met 91.

The ZIKV envelope protein (ZIKV_E) is responsible for forming the viral envelope and facilitating the virus’s entry into the target cell. It is also a primary target of neutralizing antibodies against ZIKV. Structurally, the E protein consists of three domains. Domain 1 (DI) is a central domain that connects domains 2 (DII) and 3 (DIII), and in other Flaviviruses, DI is also involved in the conformational changes necessary for viral entry into the host cell. Domain 2 (DII) is an elongated cylindrical domain containing a conserved fusion loop that interacts with the endosomal membrane of the host cell, which is essential for the release of the viral genome into the cellular cytoplasm. Finally, domain 3 (DIII) is involved in binding to cellular receptors to facilitate viral entry. In a brief structural analysis of the different domains of the E protein, domain 1 contains approximately 130 residues divided into three segments, residues 1–51, 132– 192, and 280–295. The core of domain 1 is formed into a beta-barrel structure, linked to an N-terminal filament. This type of conformation provides hydrophobic characteristics, thus decreasing its absolute solvent accessibility in this region. As observed in the experimental method used in this study, this domain did not show high membrane reactivity or high epitope probability scores. In addition to this hydrophobic characteristic, this domain also has a lack of electron density at a loop site with 150 residues, indicating a potential local glycosylation site. Domain 2 is responsible for the dimerization of the E protein, leading to an extended but interrupted dimer interface, creating voids in the dimer on both sides of domain 2. The central interface of the dimers is composed of an alpha-beta helix and J-chain elements in each monomer, while the distal dimer interface is primarily created by the hydrophobic structure of domain 1, which interacts with domains 2 and 3. Domain 3, comprising residues 296-403, displays a fold where the alpha-beta sheets and the disordered D-chain come into contact due to the loop of the adjacent E protein monomer. This loop, known as the fusion loop, is located between residues 98–109 of domain 3 of the E protein. This structure also functions in the fusion of the viral membrane with the host cell membrane during the viral entry phase. The segment where this loop is situated is highly conserved among Flaviviruses, and the most plausible hypothesis for this conservation is that this segment has an indispensable function in viral infection.

Comparatively, the envelope proteins of ZIKV, YF, USUV, WNV and SLEV are very similar regarding amino acid composition, which is quickly evident in the phylogeny. The modeled structures also show great proximity in terms of RMSD (Figure 3 - Panel A) and the PCA comparison can successfully separate them from this perspective. By analyzing the configuration of the structure via concavity, we were able to identify three putative regions for testing compounds with docking. We divided these situations into three distinct docking problems and ran the analyses as previously described. For problem 3, which returned plausible results, we used the configuration 80 80 60 for npts and the gridcenter was located at xyz coordinates -8.415 24.329 13.312, then centering in the border region of DI (190 - 193), DII (265 - 269) and finally DIII (418 - 498). These regions compared in the correlation matrices generated by Bio3D show structural similarities when analyzed pairwise (pairwise matrix figures), which in turn suggests that the pocket found is shared by these viruses in this region. We then evaluated the electrostatic charges of these viruses for this protein, specifically focusing on this region used for docking to identify substantial differences in the profiles. We observed that despite the sequence and structure similarities, the charges of the regions predicted as potential pockets for ligands are very similar between ZIKV, YF, and USUV (electronegatively charged region) but change substantially for WNV and SLEV (electropositive portions with a different signature from the three most common flaviviruses). (Figure 3 – Panels A:E). Therefore, it was necessary to rerun docking problem 3 for this region in the WNV and SLEV viruses.

**Figure 3.**
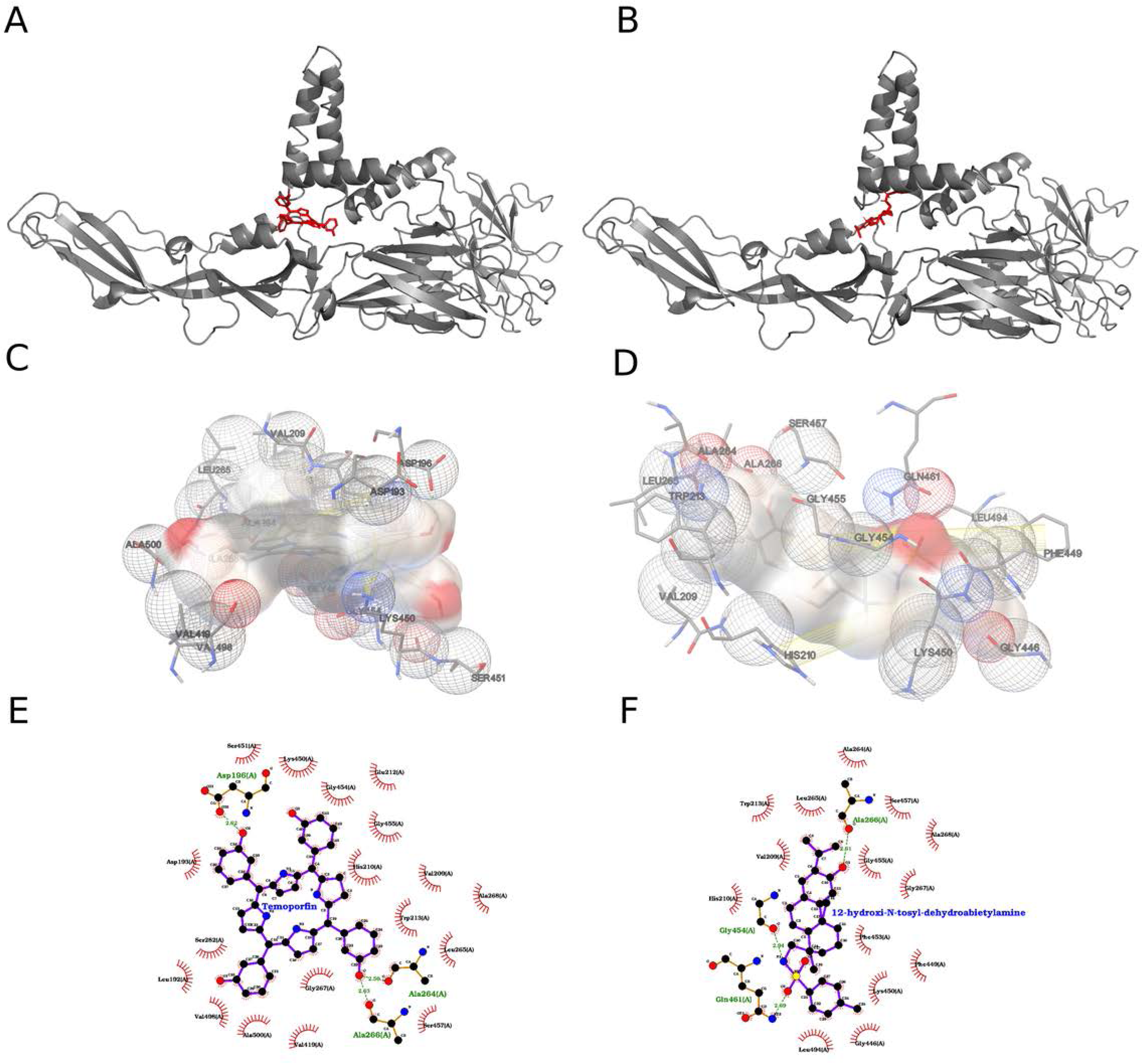
Integrated analysis of conformational ensembles across five flavivirus membrane-protein cores. (A) Two-dimensional PCA of Cα coordinates projected onto PC1 (49.9 % variance) and PC2 (35.7 %), with each 50-ps snapshot colored by virus: ZIKV (magenta), SLEV (red), USUV (blue), WNV (green), YFV (yellow). SLEV and WNV overlap in large-scale motions, YFV occupies a distinct conformer space, and ZIKV diverges most from the others. (B) Pairwise PCA projections—PC1 vs. PC2 (top left), PC2 vs. PC3 (top right), PC1 vs. PC3 (bottom left)—plus scree plot (bottom right), showing PC3 accounts for 8.7 % variance and all higher PCs contribute <1 %, thus the first three components capture the bulk of system fluctuations. (C) Ellipsoidal volume versus core size reveals a steep decay in volume as the number of core residues decreases: compact folding nuclei (<100 residues, ticks colored by virus) contrast with expansive, flexible envelopes (black bars). (D) Hierarchical clustering dendrogram (Cα-RMSD) groups SLEV (red) and YFV (yellow) most tightly (∼1 Å), then USUV (blue, ∼1.4 Å), WNV (green, ∼1.8 Å), and ZIKV (magenta, >2 Å), recapitulating PCA-derived separations. (E) Overlay of representative Cα traces highlights a conserved central α-helical scaffold (RMSD <0.5 Å) alongside variable loops/β-hairpins (up to ∼2 Å), with SLEV (red) and YFV (yellow) showing greatest congruence and ZIKV (magenta) exhibiting the largest peripheral deviation.

The interactions formed by the two compounds that gave the best results in the search with the 2000 runs for each of the 40 ligands are shown in Figure 4. In this ranking the compound temoporfin interacted with the chosen binding pocket with an energy of -12.17 kcal/mol according to the AutoDock ranking function, which translates to a *Kd* value on the order of 1.20 nM, while the compound 12-hydroxy-*N*-tosyl-dehydroabietylamine interacted with -9.78 kcal/mol and *Kd* of 68.04 nM.

**Figure 4:**
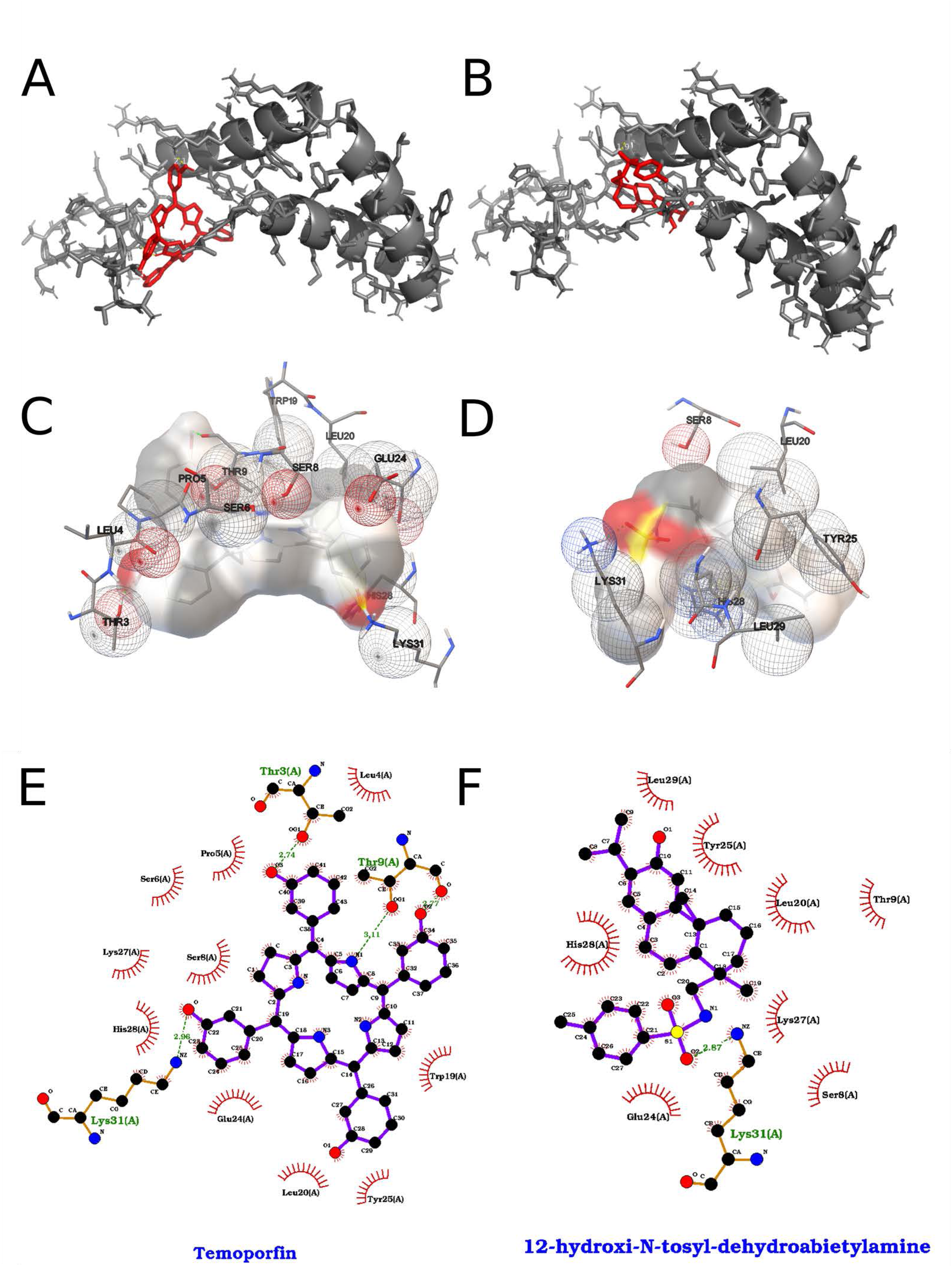
Analysis of Temoporfin and 12-hydroxy-N-tosyl-dehydroabietylamine in the target receptor pocket. (A) Ribbon representation of the receptor (grey) showing the top-scoring Temoporfin pose (red sticks) nestled into a deep cleft formed by two α-helices and an adjacent β-sheet. The chlorin ring of Temoporfin is oriented toward the back of the pocket, maximizing van der Waals contacts with the helical walls. (B) Equivalent view for the 12-hydroxy-N-tosyl-dehydroabietylamine pose (red sticks), which extends longitudinally along the groove, engaging a network of hydrogen bonds at its hydroxyl terminus. (C) Three-dimensional electrostatic surface of the pocket (semi-transparent beige) with Temoporfin bound. Key residues are shown as mesh spheres colored by interaction type: red for polar/charged, green for hydrophobic, and white for neutral. Temoporfin’s chlorin core sits adjacent to hydrophobic side chains Leu286, Val209, and Leu450, while its propionic acid side chain forms a hydrogen bond network with Asp193 (2.6 Å) and Ser198 (2.9 Å). (D) 3D surface view for 12-hydroxy-N-tosyl-dehydroabietylamine highlighting a complementary fit: the tosyl moiety occupies a hydrophobic pocket lined by Phe449, Leu494, and Ala266 (mesh spheres), whereas the hydroxy-amine end donates hydrogen bonds to Gly454 (2.7 Å) and Gln461 (2.8 Å). (E) Two-dimensional interaction diagram for Temoporfin: hydrogen bonds to Asp193 and Ser198 are shown as green dashed lines with distances (Å), π–π stacking against Phe84 is indicated by parallel ring symbols, and hydrophobic contacts (Leu286, Val209, Leu450, Ala500) are denoted by red semicircles; polar/charged residues are labeled in green. (F) Two-dimensional map for 12-hydroxy-N-tosyl-dehydroabietylamine using the same conventions: a bifurcated hydrogen-bond network to Gly454 and Gln461 (green dashed lines), extensive hydrophobic contacts (Phe449, Leu388, Leu419, Trp213) shown as red arcs, and the tosyl phenyl ring engaging Phe53 in a π–π interaction.

Temoporfin, in its best-obtained pose, formed face-to-face π-π interactions between two of its pyrrole rings and His 210 of the ZIKV_E and between one of its phenolic rings and Trp 213. Lys 450 formed a π-cation interaction with another one of the phenolic rings of temoporfin. The compound’s phenolic hydroxyl groups formed numerous hydrogen bonds with the envelope protein, specifically with the residues Asp 196, His 210, Ala 264, Lys 450, Ser 457 and Ala 500. Temoporfin formed many hydrophobic contacts with the protein, namely with the residues Leu 192, Asp 193, Val 209, His 210, Trp 213, Leu 265, Ala 268, Val 419, Val 498 and Lys 450. 12-hydroxy-*N*-tosyl-dehydroabietylamine, in its best obtained pose, formed hydrogen bonds between its sulfonamide moiety and the residues Gly 454 and Gln 461 of the ZIKV_E. Hydrogen bonding was also observed between its phenolic hydroxyl group and the residues Gly 455 and Ser 457. Due to the hydrophobic nature of the compound, several non-polar contacts were observed with the protein, namely with the residues Val 209, His 210, Trp 213, Leu 265, Ala 268, Phe 449, Lys 450 and Leu 494.

### Membrane protein

The M protein of Zika virus is a small membrane protein crucial for virus assembly and maturation. It is a type I transmembrane protein with a short N-terminal ectodomain (residues 92-130) and a C-terminal transmembrane anchor region (residues 131-166), with the ectodomain exposed on the viral membrane surface and the C-terminus embedded in the lipid bilayer. Secondary structure predictions suggest that the ectodomain is predominantly α-helical, and the transmembrane region contains two α-helical segments separated by a short loop. Experimental evidence indicates that the M protein oligomerizes, likely forming pentameric channels in the viral membrane, mediated by interactions between the transmembrane helices. During virus maturation, the M protein undergoes a conformational change from the immature precursor form (prM) to the mature M protein, triggered by the cleavage of the prM protein by the host protease furin, allowing the M protein to adopt its final arrangement in the mature virion. In summary, the M protein has a simple secondary structure consisting of an α-helical ectodomain, a transmembrane region with two membrane-spanning helices, and the ability to oligomerize into pentameric channels, undergoing a key conformational change during virus maturation essential for producing infectious virions. The M protein of dengue virus oligomerizes into a pentameric channel through structural and biochemical interactions. This process involves the M protein’s two transmembrane helices, which span the viral membrane and are crucial for oligomerization. The helices interact to form a stable pentameric channel, with specific residues such as Ala 94, Leu 95, Ser 112, Glu 124, and Phe 155 playing critical roles in stabilizing the structure. Molecular dynamics simulations have shown that the M protein forms a stable pentameric channel in an implicit membrane environment, providing insights into its structural dynamics. The pentameric channel facilitates viral release from host cells by allowing ions to pass through the membrane, a process essential for the virus life cycle and a key target for therapeutic interventions. In summary, the M protein oligomerizes into a pentameric channel through transmembrane helix interactions, specific residue mediation, and molecular dynamics simulations, playing a crucial role in viral release and presenting a target for therapeutic development. The key amino acids involved in the oligomerization of the M protein from dengue virus to form a pentameric channel include Ala 94, Leu 95, Ser 112, Glu 124, and Phe 155, which act as central hub residues that stabilize the pentameric structure through physicochemical interactions between the transmembrane domains. Additionally, amino acids such as Glu, Thr, Ser, Trp, Ala, and Ile form the pore-lining residues of the channel, conferring an overall negative charge. Hydrophilic amino acids located near the symmetry axis of the oligomer and between subunits contribute to the distortion from perfect symmetry. Regions modulating oligomerization contain a higher fraction of polar, charged residues, glycine, and proline compared to conventional interfaces and protein surfaces, mediating specific interactions and preventing nonspecific dysfunctional aggregation. In summary, specific residues like Ala 94, Leu 95, Ser 112, Glu 124, and Phe 155 are crucial for oligomerization, with pore-lining residues being predominantly hydrophilic, and regions modulating oligomerization enriched in polar, charged residues, glycine, and proline. Hydrophilic amino acids play a crucial role in modulating the oligomerization of the M protein from dengue virus into a pentameric channel structure. These amino acids form the pore-lining residues of the pentameric channel, conferring an overall negative charge. Located near the symmetry axis of the oligomer and between subunits, they contribute to the distortion from perfect symmetry. Regions modulating oligomerization contain a larger fraction of polar, charged residues compared to conventional interfaces and protein surfaces, mediating specific interactions and preventing nonspecific dysfunctional aggregation. Despite this, there is no significant change in the ratio of hydrophilic to hydrophobic amino acids between different conformers of the M protein, indicating that the main source of structural readjustment is due to multiple polar asymmetric interactions of hydrogen bonds. In summary, hydrophilic amino acids in the M protein are essential for oligomerization by forming pore-lining residues, contributing to symmetry distortion, mediating specific interactions, and maintaining the structural integrity of the pentameric channel. Hydrophilic and hydrophobic amino acids play crucial roles in the oligomerization of the M protein from dengue virus into a pentameric channel structure. The pore-lining residues, predominantly hydrophilic amino acids such as glutamic acid (Glu), threonine (Thr), serine (Ser), tryptophan (Trp), alanine (Ala), and isoleucine (Ile), confer an overall negative charge to the channel pore. These hydrophilic residues, located near the symmetry axis of the oligomer and between subunits, contribute to symmetry distortion and stabilize the pentameric structure through specific hydrogen bonding and polar interactions. Regions modulating oligomerization contain a larger fraction of polar, charged residues, glycine, and proline compared to conventional protein interfaces, which mediate specific interactions and prevent nonspecific dysfunctional aggregation of the M protein. Additionally, key hydrophobic amino acids, such as alanine (Ala 94), leucine (Leu 95), and phenylalanine (Phe 155), stabilize the transmembrane helix interactions and the overall pentameric channel structure. In summary, hydrophilic amino acids in the M protein are essential for forming pore-lining residues, contributing to symmetry distortion, mediating specific interactions, and preventing nonspecific aggregation, while working in concert with hydrophobic amino acids to stabilize the pentameric channel structure.

Figure 5 shows the results for comparing the modeled structures for the five viruses regarding the structural protein M. A 2D PCA projection (PC1 vs. PC2) explains 48.56% and 21.3% of the variance, respectively, revealing a clear structural divergence among the viruses, with YFV and ZIKV being the most distant and WNV and USUV closely grouped, supporting their phylogenetic proximity (Panel A). This perspective is expanded with additional projections and a scree plot indicating that the first three principal components capture over 86% of the structural variance, validating the observed clusters (Panel B). The volumetric profile of the protein’s core region increases sharply from ∼65 residues, suggesting greater structural variability, while red bars indicate conserved structural cores critical for comparative analyses (Panel C). Hierarchical clustering reveals two main groups: one with WNV and USUV, and another with ZIKV and SLEV, while YFV diverges early, reflecting its distinct evolutionary lineage (Panel D). Structural superposition highlights highly conserved α-helices in the upper region and variability in loop regions, particularly in ZIKV and YFV, suggesting potential functional adaptations or antigenic differences. WNV and USUV exhibit nearly identical alignments in β-sheets and helices, confirming their structural closeness (Panel E). Multiple sequence alignment reinforces conservation in internal structural regions (helices and sheets) and variations in exposed areas, especially between ZIKV and YFV, pointing to divergence possibly related to self assembly mechanisms not discussed here, while WNV and USUV show high similarity (Panel F). These analyses highlight the marked structural and evolutionary divergence of YFV, the close relationship between WNV and USUV, and the intermediate position of SLEV. They also identify conserved structural cores crucial for developing broad-spectrum vaccines or drugs which were then explored in futher analyses herein.

**Figure 5.**
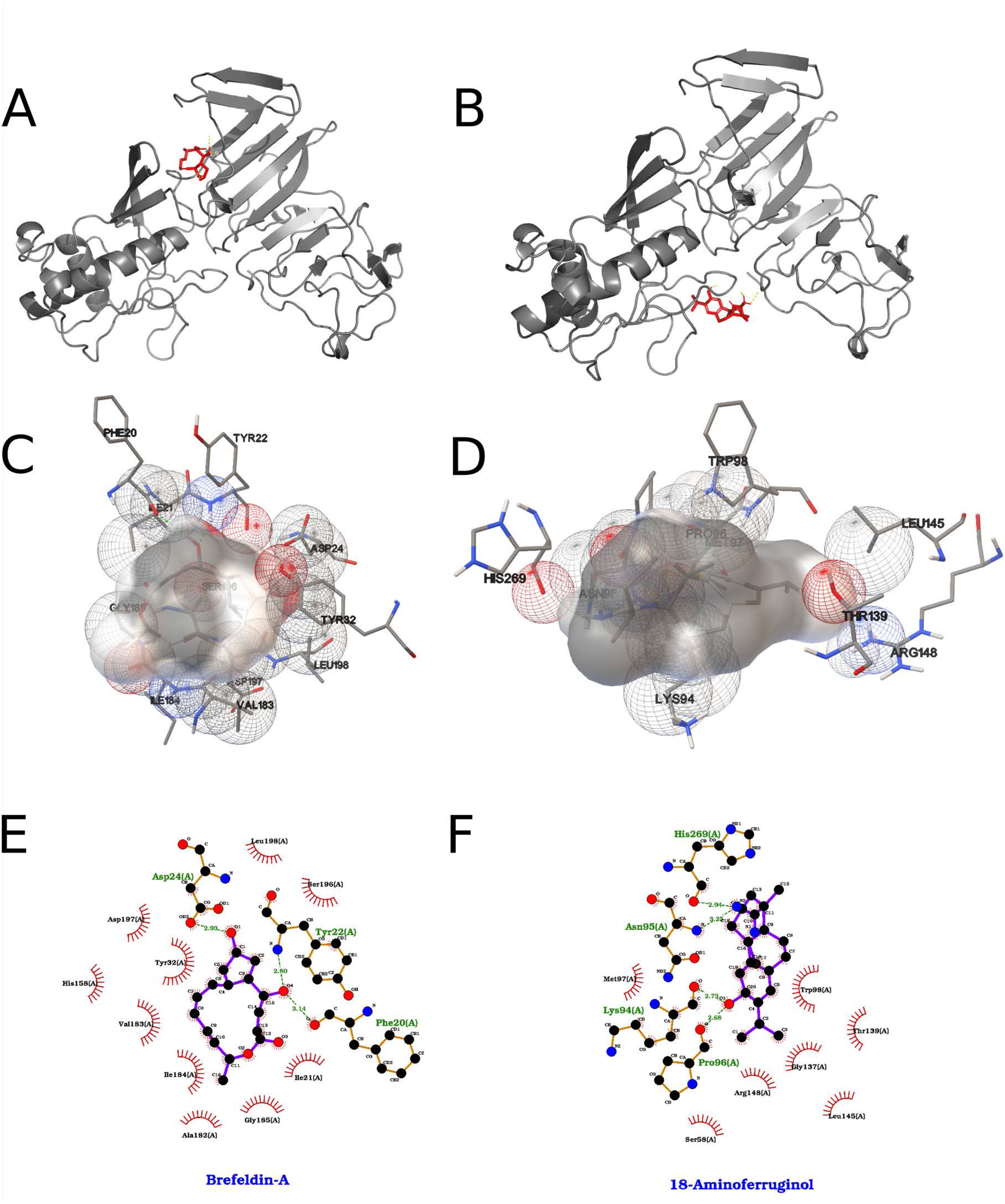
Conformational and sequence-based comparison of the flavivirus membrane protein core. (A) Two-dimensional principal component analysis (PCA) of the Cα-atom coordinates from molecular dynamics trajectories, projected onto the first two principal axes (PC1 vs. PC2). PC1 accounts for 48.6 % of the total variance and separates the five virus homologs along the horizontal axis; PC2 accounts for 21.3 % of the variance on the vertical axis. Each point represents one 50-ps snapshot (n = 200 per system) colored by virus: ZIKV (magenta), SLEV (red), USUV (blue), WNV (green), YFV (yellow). (B) Pairwise projections of the same snapshots onto PC1 vs. PC2 (top left), PC1 vs. PC3 (top right), and PC2 vs. PC3 (bottom left), illustrating that PC3 (17.1 % variance) captures additional orthogonal motions. Bottom right: scree plot of eigenvalue rank vs. proportion of variance explained, highlighting that the first three components together cover >86 % of system fluctuations (48.6 % + 21.3 % + 17.1 % = 86.9 %). (C) Core-size vs. total ellipsoidal volume analysis. For each structure, the subset of residues forming the most tightly packed core was defined by iteratively removing the most mobile residues until the remaining Cα ensemble occupied <10 Å³ of ellipsoidal volume; the x-axis shows the number of residues in that core, the y-axis the corresponding ellipsoidal volume. The five homologs delineate descending curves; the smallest, most compact core (∼55 residues) is observed in ZIKV (yellow tick), whereas YFV (magenta tick) retains a larger core (∼65 residues). (D) Hierarchical clustering of the five equilibrated structures based on pairwise backbone RMSD (Å) distances. The dendrogram reveals two major clusters: {ZIKV, SLEV} vs.{WNV, USUV, YFV}, with ZIKV (yellow) and SLEV (red) more similar to each other (RMSD ∼1.2 Å) than to the others. (E) Structural superposition of the five representative equilibrated cores, shown as ribbons. Coloring follows the same virus scheme: ZIKV (yellow), SLEV (red), USUV (blue), WNV (green), YFV (magenta). The overlay highlights that the long central α-helix is highly conserved in orientation and length, whereas the distal loop region exhibits the greatest divergence in backbone trace. (F) Sequence alignment heatmap of the core region (residues 100–170) colored by residue frequency across the five viruses (blue = highly conserved; red/yellow = variable). The central α-helix (positions 120–150) shows >90 % identity (predominantly polar and hydrophobic heptad repeats), whereas loops flanking the helix display scattered substitutions (notably at positions 105, 115, 155).

The interactions formed by the two compounds that gave the best results in the search with the 2000 runs for each of the 40 ligands are shown in Figure 6. In this ranking the compound temoporfin interacted with the chosen binding pocket with an energy of -8.59 kcal/mol according to the AutoDock ranking function, which translates to a *Kd* value on the order of 503.88 nM, while the compound 12-hydroxy-*N*-tosyl-dehydroabietylamine interacted with -7.39 kcal/mol and *Kd* of 3.84 µM.

**Figure 6.**
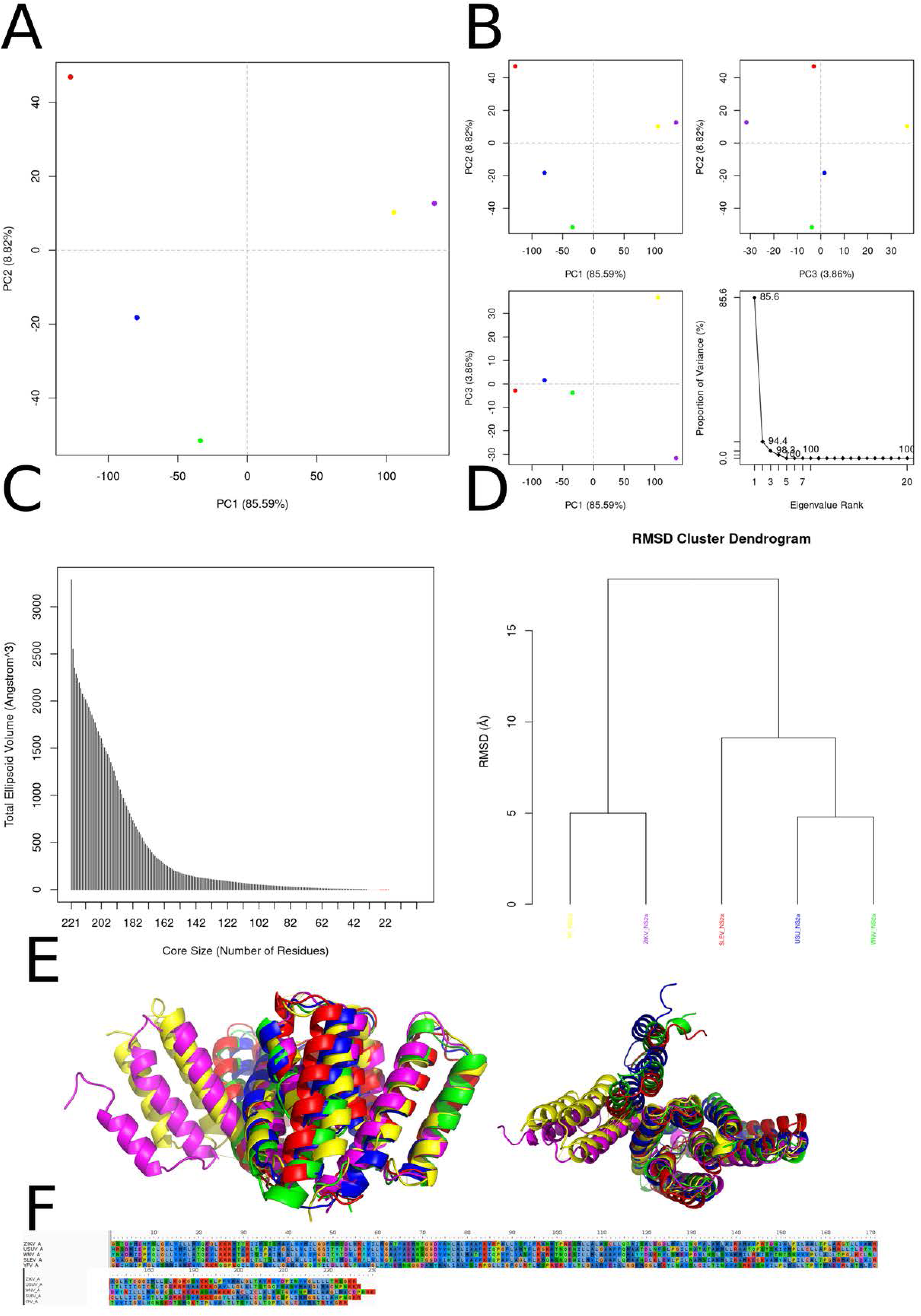
Best results for Temoporfin and 12-hydroxy-N-tosyl-dehydroabietylamine dockings againts Membrane protein, where on the left we represent Temoporfin and on the right 12-hydroxy-N-tosyl-dehydroabietylamine. (A, B) Orthogonal three-dimensional views of the ligand (highlighted in red sticks) positioned within the target protein’s binding site (gray cartoon and stick representation). (C, D) Molecular surface rendering of the pocket colored by electrostatic potential (white-to-red scale for negative regions, white-to-blue for positive), with neighboring side chains shown as transparent spheres to emphasize the binding cavity. (E, F) Two-dimensional interaction diagrams (LigPlot-style), depicting hydrogen bonds as green dashed lines with bond distances (Å) and hydrophobic contacts as red semicircles around the relevant ligand atoms. This layout is repeated for each of the seven screened compounds in their respective protein pockets.

Temoporfin, in its best-obtained pose, formed π-π interactions of the face-to-edge type between two of its phenolic rings and one of its pyrrole rings and Trp 19 of the ZIKV_M and between one of these two phenolic rings and Tyr 25 of the protein. One of the other pyrrole and phenolic rings of temoporfin formed π-π interactions of the face-to-face and face-to-edge, respectively, with the His 28 of the ZIKV_M. The compound formed hydrogen bonds between its phenol groups and the side chain of Thr 3, the main chain carbonyl oxygen of Thr 9 and the side chain of Lys 31. One of the compound’s pyrrole nitrogen atom formed a hydrogen bond with the side chain of the residue Thr 9. The compound formed several hydrophobic contacts with the protein, namely with the residues Thr 3, Pro 5, Ser 6, The 9, Leu 20, Glu 24 and Lys 27.

12-hydroxy-*N*-tosyl-dehydroabietylamine, in its best obtained pose, formed hydrogen bonds between its sulfonamide moiety and the residues His 28 and Lys 31 of the ZIKV_M. The compound formed π-π interactions of the fate-to-edge type between its toluene ring and the residue His 28 of the protein, of the edge-to-edge kind between its phenolic ring and Tyr 25 and of the face-to-face type between the phenolic ring and the residue His 28. Due to the hydrophobic nature of the compound, several non-polar contacts were observed with the protein, namely with the residues Thr 9, Trp 19, Leu 20, Glu 24, Tyr 25, His 28, Leu 29, Lys 31, Val 32, Pro 72 and Ala 73.

### NS1 Protein

The non-structural protein 1 (NS1) plays a multifactorial virulence function in Flavivirus. Preliminary data are also under analysis for other structural and non-structural Flavivirus proteins. NS1 is highly conserved among flaviviruses, containing 12 cysteines and being glycosylated similarly to other non-structural viral proteins. Glycosylated NS1 associates with lipids to form a homodimer within cells, which is necessary for late-stage viral replication. Additionally, NS1 is secreted into the extracellular space as a hexameric lipoprotein particle involved in immune evasion and pathogenesis through interactions with both innate and adaptive immune system components, as well as other host factors. NS1 is a primary antigenic marker of viral infection and is used alongside other markers for early detection of DENV infection. The NS1 coding sequence is suspected to be a key genetic factor underlying the diverse clinical outcomes of infections caused by more than 70 flavivirus members. However, ZIKV NS1 appears to have a different pathogenic relationship compared to other flaviviruses. Flavivirus NS1 protein has three distinct domains: domain one, an N-terminal β-barrel (“β-roll”) hydrophobic region from amino acid 1 to 29; domain two, an α/β wing from amino acids 38 to 151; and a C-terminal β-ladder central region with an extensive β-sheet from amino acids 181 to 352. It has twelve invariant cysteines forming six disulfide bonds per monomer. The segments between these domains (30-37 and 152-180) form a “connector” subdomain, comprising 3 β-sheet chains linking the wing to the central β-barrel and β-ladder domains. The fundamental unit is a cross-shaped flat dimer formed by end-to-end β-ladder meshes. The inner face of the dimer, comprising the β-barrel domain and adjacent loop, forms a hydrophobic surface. The outer face of the dimer is polar and contains glycosylation sites. In the NS1 hexamer, three dimers join with the polar glycosylated faces outward and the hydrophobic faces inward, enabling interaction with lipid molecules in the secreted NS1 lipoprotein particle.

The principal component analysis (PCA) of the NS1 “wing” domain reveals a highly structured conformational landscape in which the first three modes capture nearly 88 % of the total variance. Along PC1 (52.65 %), USUV and ZIKV cluster tightly together, separated from WNV, SLEV and YFV, suggesting that a dominant loop-oriented motion distinguishes these two species from the others. PC2 (22.01 %) further discriminates YFV from the WNV/SLEV pair, implicating an orthogonal displacement—likely a subtle β-sheet tilt or loop swing—unique to YFV. Even PC3 (13.23 %) contributes appreciably to the spread of WNV and SLEV conformations, indicating that their wing domains sample additional collective motions not accessible to USUV/ZIKV, whose PC3 dispersion remains minimal (Figure 7 – Panel A). To define a robust structural core, we calculated the minimal enclosing ellipsoid volume over progressively smaller residue sets. As peripheral residues were trimmed, the ellipsoidal volume rapidly declined before stabilizing in a plateau region (∼200–250 residues) and then dropping sharply to a minimum at approximately 150–170 residues. This inflection identifies a tightly concerted scaffold—composed principally of β-strands and connecting turns—whose dynamics underlie the domain’s essential fold, while excluding highly mobile loops that inflate the overall volume. Selecting roughly 160 residues thus balances retention of key secondary-structure elements with removal of peripheral fluctuations (Figure 7 – Panel B). Hierarchical clustering by pairwise RMSD of these core centroids mirrors both the PCA grouping and known phylogenetic relationships. USUV and ZIKV exhibit the smallest mutual RMSD (<0.4 Å), reflecting nearly identical backbone conformations. WNV shows intermediate divergence (∼0.7 Å), whereas SLEV and YFV branch off with larger deviations (∼0.9 Å and >1.1 Å, respectively). These modest Ångström-scale differences confirm global conservation of the wing fold, with species-specific nuances arising from slight reorientations of loops rather than wholesale topological changes (Figure 7 – Panel C). A structural overlay of the six centroids accentuates this picture: the β-sheet core aligns within ∼0.5 Å across all species, while surface-exposed loop regions fan out by up to 3–4 Å. Such loops mediate interactions in NS1 hexamers and with host factors; their variability likely tunes viral assembly, immune recognition and host specificity. Indeed, the compact loops of USUV/ZIKV align with their tight PCA clustering, whereas the more dispersed loops of WNV, SLEV and YFV reflect broader conformational sampling. Corresponding multiple-sequence alignments show that conserved hydrophobic residues anchor the β-strands, while charged or polar substitutions cluster in the flexible loops (Figure 7 – Panel D). Together, these findings link sequence divergence to dynamic behavior, suggesting that evolution has preserved the core scaffold of NS1 while permitting loop flexibility to drive species-specific functions and immunological profiles. 3D structural superposition (Figure 7 -Panel E) highlights high conservation in the β-ladder and central β-sheet across all viruses, particularly between WNV and USUV, whereas ZIKV and SLEV exhibit loop and terminal helix variations, and YFV displays pronounced deviations in the wing domain and dimer interfaces, likely influencing host interactions and immune evasion. MSA (Panel F) further corroborates these findings, showing strong sequence conservation in internal β-sheets but high variability in surface-exposed loops and glycosylation sites, with WNV and USUV nearly identical, ZIKV and SLEV sharing key motifs but differing in insertions, and YFV exhibiting the most sequence divergence, even in typically conserved regions. Collectively, these analyses underscore YFV’s unique structural and evolutionary trajectory, the close structural relationship between WNV and USUV, and the intermediate divergence of ZIKV and SLEV, while identifying a conserved structural core (120– 160 residues) as a potential target for broad-spectrum antivirals. This integrated approach advances our understanding of NS1’s structure-function relationships and provided a framework for our screening process.

**Figure 7:**
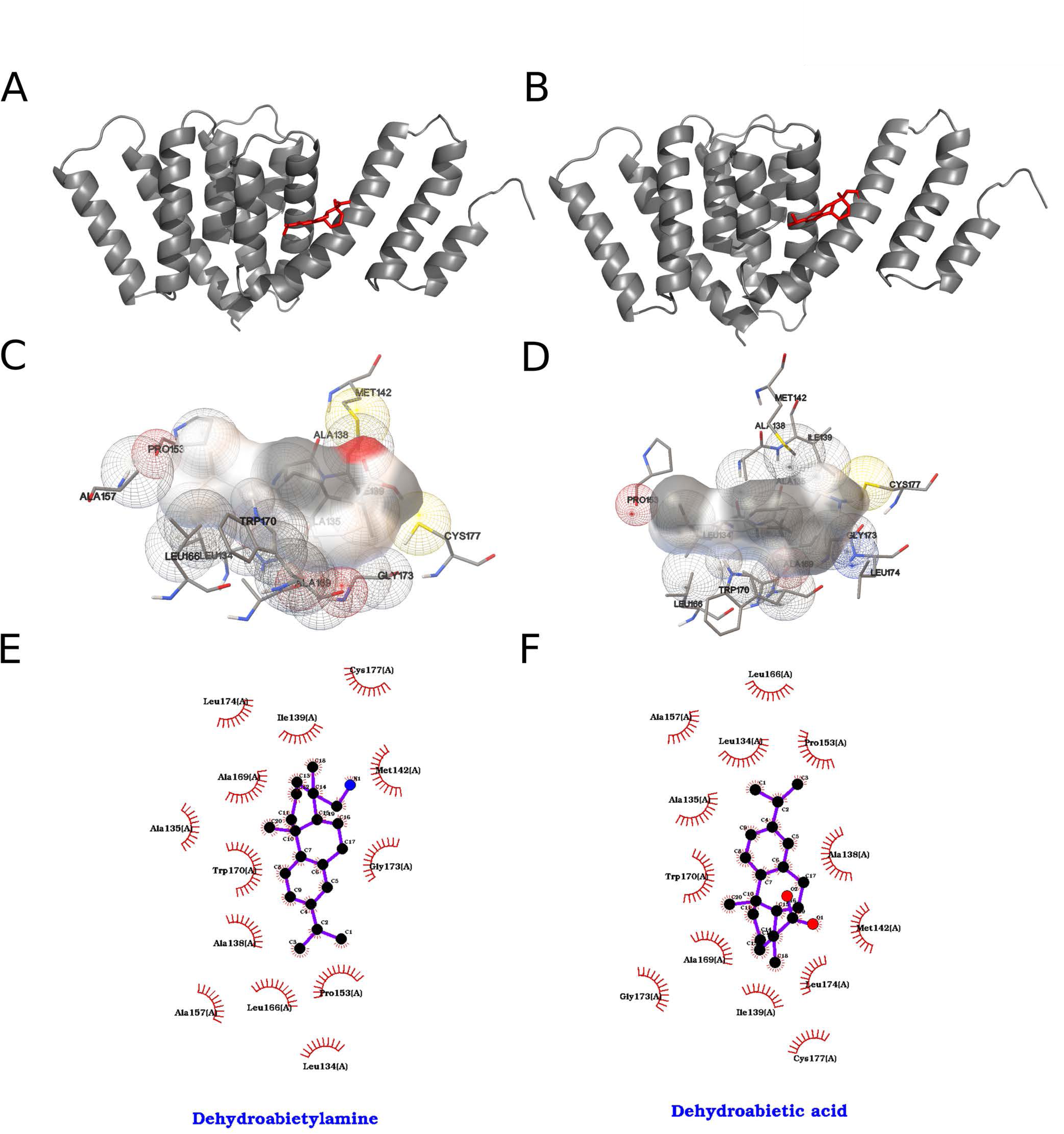
(A) Two-dimensional projection of the central conformations from six molecular dynamics simulations of the NS1 “wing” domain for five flavivirus species in principal component space. Each colored dot represents the centroid of a conformational cluster: ZIKV (magenta), USUV (blue), WNV (green), SLEV (red), and YFV (yellow). The axes correspond to PC1 (52.65 % of total variance) and PC2 (22.01 %). Dispersion along PC1 highlights the primary conformational differences among the variants. (B) Complementary PCA plots and variance scree: (top left) PC1 vs. PC2; (top right) PC2 vs. PC3 (13.23 % of variance); (bottom left) PC3 vs. PC1; (bottom right) scree plot showing the cumulative variance explained by each eigenvalue, with a steep drop after the first three components. (C) Total ellipsoidal volume of the structural “core” — defined as the minimal enclosing ellipsoid of the “wing” domain — plotted against the number of residues included. Bars indicate that as the core size decreases, the volume plateaus (light pink region) before reaching a minimum at approximately 150–170 residues (red), suggesting an optimal core definition. (D) Hierarchical clustering dendrogram based on pairwise RMSD between the central conformations. Branch heights represent the average Ångström deviation required to align cluster centroids. USUV (blue) and ZIKV (magenta) cluster together, indicating high conformational similarity, whereas WNV (green), SLEV (red), and YFV (yellow) form distinct branches. (E) Structural overlay of the six cluster centroids of the NS1 “wing” domain, rendered as ribbon traces and color-coded as in panels A–D. The close alignment of β-sheets underscores secondary-structure conservation, while loop regions display greater variability. (F) Multiple sequence alignment of the amino-acid segments corresponding to the NS1 “wing” domain for the five species, shown in three blocks (residues 1–50, 51–100, and 101–150). Background coloring denotes physico-chemical residue properties (hydrophobic, polar, charged), highlighting strongly conserved positions and surface-exposed variable hotspots, particularly within loop regions.

The interactions formed by the two compounds that gave the best results in the search with the 2000 runs for each of the 40 ligands are shown in Figure 8. In this ranking the compound 18-aminoferruginol interacted with the chosen binding pocket with an energy of -8.50 kcal/mol according to the AutoDock ranking function, which translates to a *Kd* value on the order of 583.58 nM, while the compound Brefeldin A interacted with -8.45 kcal/mol and *Kd* of 639.67 nM.

**Figure 8.**
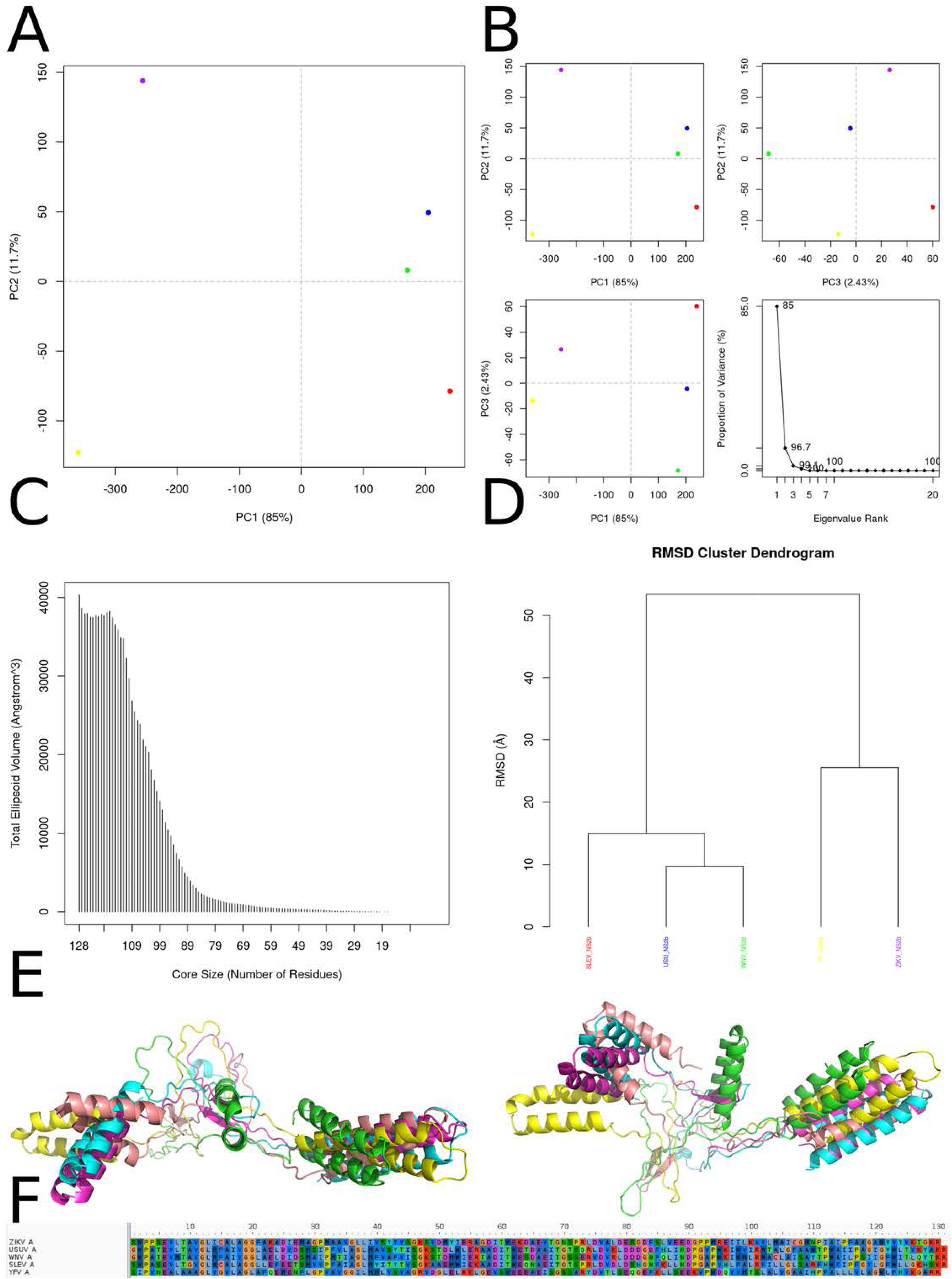
Best results for 18-aminoferruginol and Brefeldin A dockings against NS1 protein, where on the left we represent 18-aminoferruginol and on the right Brefeldin A. (A, B) Orthogonal three-dimensional views of the ligand (highlighted in red sticks) positioned within the target protein’s binding site (gray cartoon and stick representation). (C, D) Molecular surface rendering of the pocket colored by electrostatic potential (white-to-red scale for negative regions, white-to-blue for positive), with neighboring side chains shown as transparent spheres to emphasize the binding cavity. (E, F) Two-dimensional interaction diagrams (LigPlot-style), depicting hydrogen bonds as green dashed lines with bond distances (Å) and hydrophobic contacts as red semicircles around the relevant ligand atoms.

18-aminoferruginol, in its best obtained pose, formed a face-to-edge π-π interaction between its phenolic ring and Trp 98 of the ZIKV_NS1, while the phenolic hydroxyl group formed a hydrogen bond with the backbone carbonyl oxygen atom of the residue Pro 96. The compound’s amine moiety formed hydrogen bonds with the backbone carbonyl oxygen atoms of the residues Asn 95 and His 269. Given its nonpolar nature, the compound formed many hydrophobic contacts with the protein, namely with the residues Lys 94, Pro 96, Asp 136, Asp 138, Thr 139, Leu 145, Arg 148, His 269 and Val 350 (Figure 8 – A, C, E).

Brefeldin A, in its best obtained pose, formed hydrogen bonds between its cyclopentane ring hydroxyl group and the residue Asp 24 of the ZIKV_NS1 and between its other hydroxyl moiety and the backbone carbonyl oxygen atom of the residue Phe 20. The compound formed π-π interactions between the unsaturation distal to its lactone moiety and the residue Tyr 32 of the NS1 and between its other unsaturation and the residue Tyr 22. The compound formed some non-polar contacts with the protein, namely with the residues Val 19, Ile 21, His 158, Val 183 and Leu 198 (Figure 8 – B, D, F).

### NS2a protein

The NS2a protein of ZIKV is a non-structural protein that plays a crucial role in the viral life cycle, particularly in the assembly and morphogenesis of the virus. Understanding its secondary and tertiary structures is essential for elucidating its function. The secondary structure of ZIKV NS2a is characterized by the presence of multiple transmembrane (TM) domains and helical segments. Biochemical studies suggest that NS2a has a single segment that traverses the endoplasmic reticulum (ER) membrane, alongside six peripheral segments that associate with the membrane. The transmembrane domain is likely to be α-helical, which is common in membrane proteins, facilitating its integration into the lipid bilayer. The specific arrangement of these helical segments contributes to the protein’s ability to interact with other viral components. For example, NS2a is known to recruit genomic RNA and structural proteins (C, prM, and E) to the virion assembly site, indicating that its secondary structure is integral to these interactions. The tertiary structure of NS2a is less well-defined compared to its secondary structure, primarily due to its dynamic nature and interactions with the lipid environment. The protein’s tertiary conformation is influenced by its interactions with the ER membrane and other viral proteins. Structural probing indicates that NS2a’s conformation is adaptable, allowing it to participate in the assembly of the viral replication complex. The tertiary structure likely involves a combination of hydrophobic and charged residues that facilitate its anchoring to the membrane and interactions with viral RNA and proteins. This adaptability is crucial for its role in recruiting components necessary for virion assembly and ensuring efficient viral replication.

First, the principal-component analysis (PCA) in Panel A makes clear that PC1—accounting for 85.6 % of the total variance—dominates the conformational landscape of NS2a. Variants from Yellow fever virus (YFV; yellow), Zika virus (ZIKV; purple), West Nile virus (WNV; green), Usutu virus (USUV; blue), and Saint Louis encephalitis virus (SLEV; red) occupy discrete positions along PC1, indicating that the relative tilting and rotational packing of the four-helix transmembrane bundle are virus-specific signatures. These differences in helix bundle orientation likely modulate how NS2a embeds in the endoplasmic reticulum membrane and interacts with other nonstructural proteins (e.g., NS4A, NS4B) within the replication complex. PC2 (8.8 %) and PC3 (3.9 %), when plotted against PC1 or each other in Panel B, reveal additional, subtler variations primarily localized to extra membrane loops and helix termini— regions implicated in recruiting host factors such as RNA-binding proteins and membranes curvature modulators. Panel C’s ellipsoidal-volume analysis of the conformational “core” quantifies how volume declines steeply as residues are pruned from 221 down to approximately 50. Above ∼100 residues, the volume drop is pronounced, reflecting the removal of peripheral, mobile loops and short helices; below ∼50 residues, the curve flattens, indicating that the remaining segment constitutes a minimal, rigid scaffold. This undeformable core—composed of the four longest helices—is essential for preserving the topology required for paired-helix membrane insertion and for maintaining the geometry of protein–protein interfaces within the replicase. The RMSD clustering dendrogram in Panel D further underscores these observations: NS2a models segregate into two major clades (SLEV+ZIKV vs. USUV+WNV+YFV), with inter-clade distances of 10–15 Å and intra-clade distances of 4–6 Å. These divergence scales mirror known phylogenetic relationships and suggest that overall fold conservation masks discrete shifts in helix register and twist that might affect the assembly kinetics or stability of the NS2a-containing replicase organelle. Panel E’s structural superposition highlights a near-perfect alignment of the central core helices (H1–H4) across all species, while helix H5 and the two extramembrane loops adopt variable trajectories. Notably, the loop connecting H2 and H3 shifts by up to 3–4 Å between SLEV and USUV, a movement that could tune accessibility to the NS3 helicase or the host lipid-scramblase TMEM41B. Similarly, the N-terminal amphipathic region displays modest swing angles, potentially altering its membrane curvature sensing. The 50-residue core alignment in Panel F reveals complete conservation of bulky hydrophobic residues (Leu, Ile, Val) at inter-helix interfaces, forming an immobile hydrophobic spine. Interspersed glycine and proline residues occupy strategic hinge positions—introducing the flexibility needed for PC2/PC3 motions without compromising core integrity. This elegant balance between a sturdy hydrophobic core and locally adaptive hinges likely enables NS2a to maintain a universal membrane-anchored scaffold while fine-tuning its conformational ensemble for optimal replication complex formation and host adaptation in distinct flavivirus lineages.

The interactions formed by the two compounds that gave the best results in the search with the 2000 runs for each of the 40 ligands are shown in Figure 10. In this ranking the compound dehydroabietic acid interacted with the chosen binding pocket with an energy of -7.01 kcal/mol according to the AutoDock ranking function, which translates to a *Kd* value on the order of 7.30 µM, while the compound dehydroabietylamine interacted with -6.73 kcal/mol and *Kd* of 11.71 µM. In this case, the two best results were very similar between themselves.

**Figure 9.**
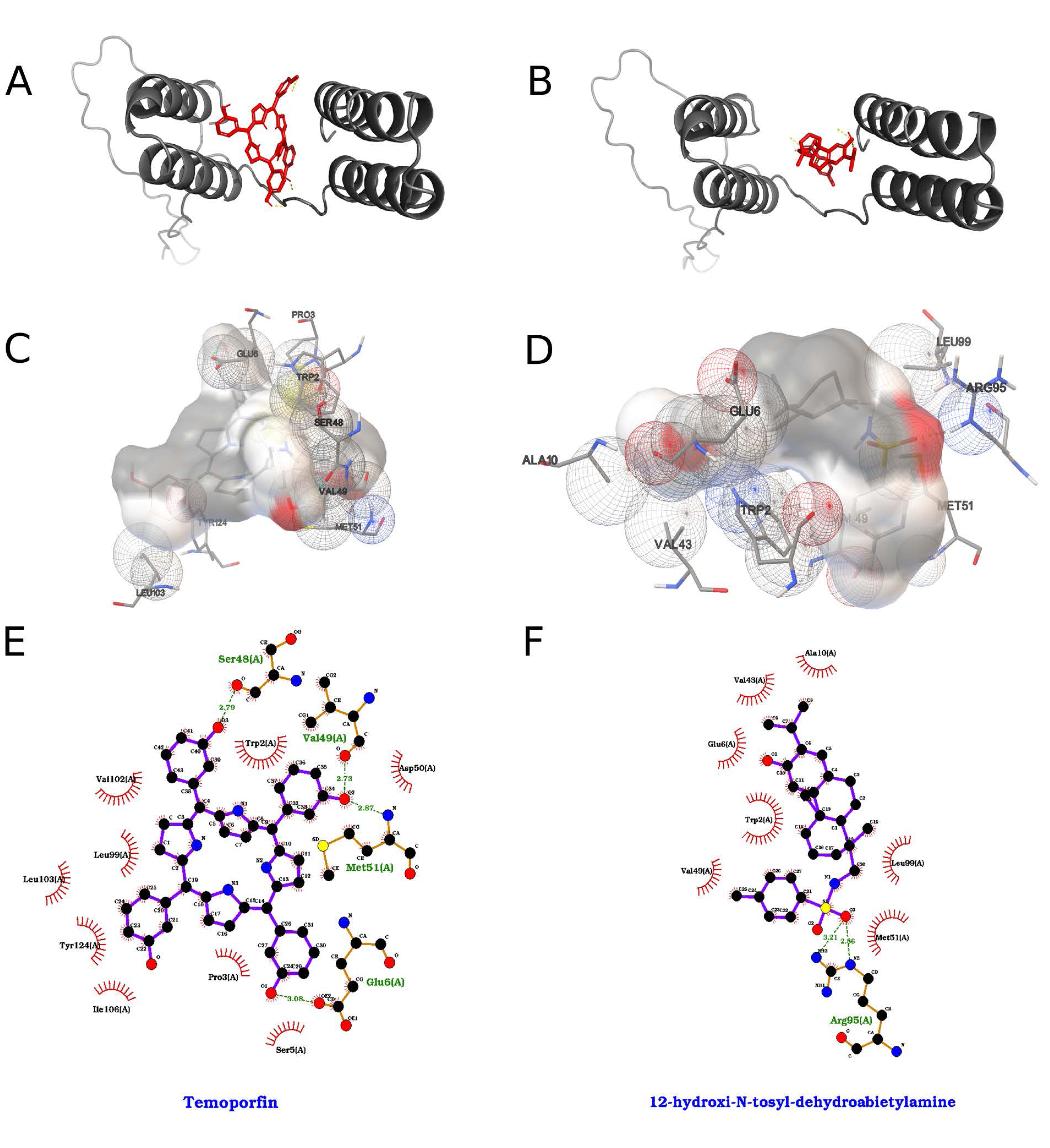
Conformational diversity, core stability, and sequence conservation of NS2a across five flaviviruses. (A) Principal-component analysis (PCA) of the NS2a conformational ensemble projected onto the first two principal axes (PC1 vs. PC2; 85.6% and 8.8% of total variance, respectively) reveals clear separation of Zika virus (magenta), West Nile virus (green), Usutu virus (blue), Saint Louis encephalitis virus (red), and Yellow fever virus (yellow) models. (B) PCA sub-projections of PC1 vs. PC2 (top left), PC1 vs. PC3 (top right; PC3 accounts for 3.9% of variance), and PC2 vs. PC3 (bottom left) emphasize that PC1 dominates the conformational differences among species. The scree plot (bottom right) confirms that over 94% of the variance is captured by the first three eigenvectors. (C) Dependence of the total ellipsoidal volume of the NS2a core on core size (number of residues). As the core is reduced from 221 to ∼50 residues, volume decreases sharply, indicating that a compact subset of helices underlies the structurally invariant scaffold. (D) Hierarchical clustering of pairwise root-mean-square deviations (RMSDs) groups NS2a models by virus species, reflecting species-specific “fingerprints” in helix packing despite overall structural homology. (E) Structural superposition of representative NS2a models (same color coding as in A) highlights a conserved four-helix bundle that mediates membrane association, overlaid by variable extramembrane loops that may confer species-specific interactions with viral and host factors. (F) Sequence alignment of the 50-residue conformational core shows invariant hydrophobic residues (shaded) at helix interfaces and conserved glycine and proline motifs that likely facilitate helix bending and packing; these conserved elements underpin the architectural integrity of the replication complex across flaviviruses.

**Figure 10.**
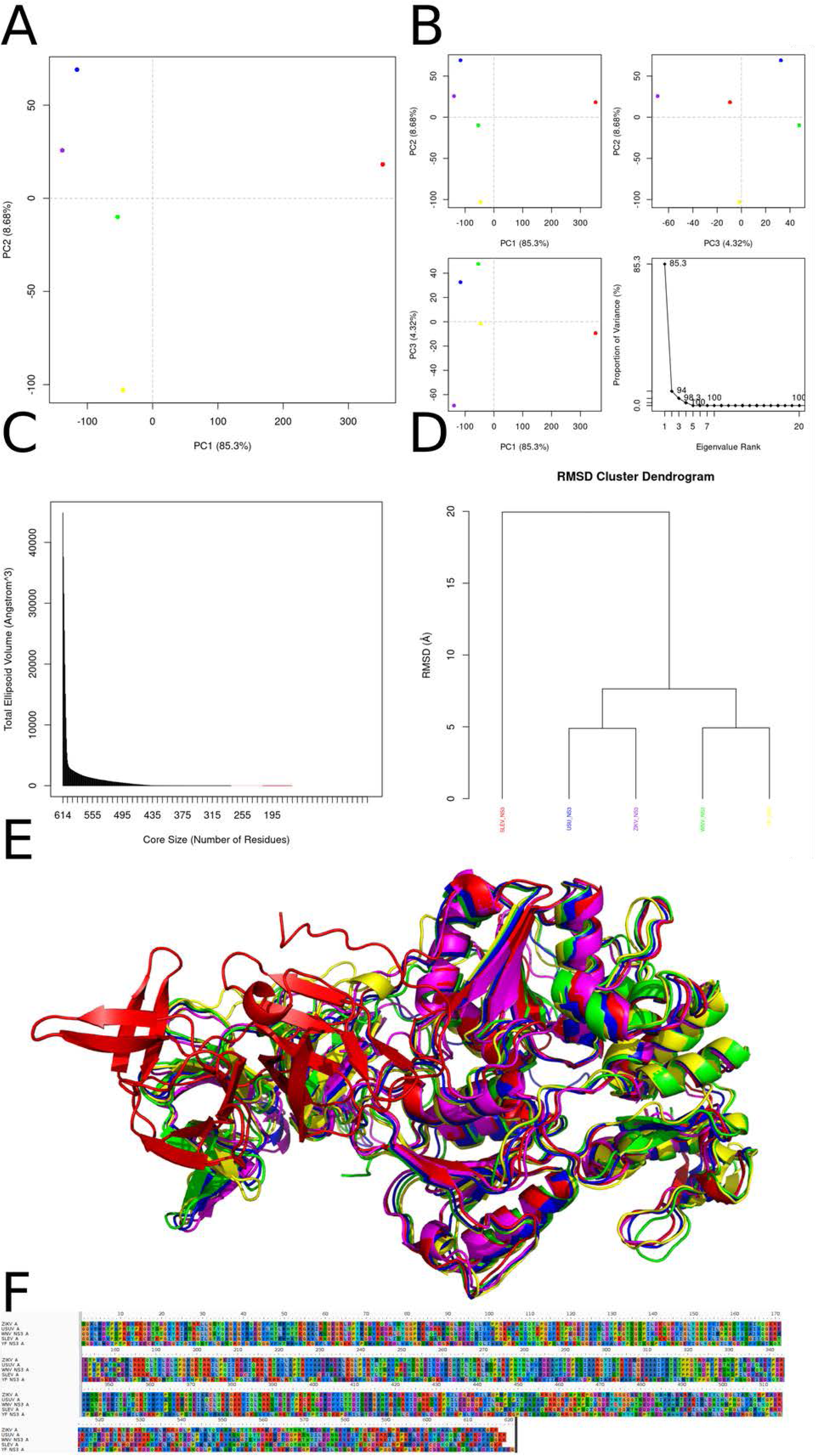
Best results for dehydroabietic acid and dehydroabietylamine dockings against NS2a protein, where on the left we represent dehydroabietic and on the right dehydroabietylamine. (A, B) Orthogonal three-dimensional views of the ligand (highlighted in red sticks) positioned within the target protein’s binding site (gray cartoon and stick representation). (C, D) Molecular surface rendering of the pocket colored by electrostatic potential (white-to-red scale for negative regions, white-to-blue for positive), with neighboring side chains shown as transparent spheres to emphasize the binding cavity. (E, F) Two-dimensional interaction diagrams (LigPlot-style), depicting hydrogen bonds as green dashed lines with bond distances (Å) and hydrophobic contacts as red semicircles around the relevant ligand atoms.

Dehydroabietic acid, in its best obtained pose, formed a non-conventional hydrogen bond between its carboxylic acid moiety and the residue Cys 177 of the ZIKV_NS2a protein. The compound formed π-π interactions of the face-to-edge type between its benzene ring and the residue Trp 170. Owing to the compounds’ hydrophobic nature, it formed several hydrophobic contacts with the protein, namely with the residues Leu 134, Ala 135, Ala 138, Ile 139, Met 142, Pro 153, Ile 154, Ala 157, Leu 166, Ala 169, Gly 173 and Leu 174.

Dehydroabietylamine, in its best obtained pose, formed a non-conventional hydrogen bond between its amine group and the residue Cys 177 of the ZIKV_NS2a protein. The compound formed π-π interactions of the face-to-edge type between its benzene ring and the residue Trp 170. Owing to the compounds hydrophobic nature, it formed several hydrophobic contacts with the protein, namely with the residues Leu 134, Ala 135, Ala 138, Ile 139, Met 142, Pro 153, Ile 154, Ala 157, Leu 166, Ala 169, Gly 173 and Leu 174.

### NS2b protein

The NS2b protein of the Zika virus (ZIKV) is a small, hydrophobic protein that acts as a cofactor for the viral NS3 protease. While the secondary and tertiary structures of the NS2b-NS3 protease complex have been studied, there is limited information specifically on the secondary and tertiary structure of the NS2b protein alone. The NS2b protein is believed to have a secondary structure consisting primarily of transmembrane helices and peripheral segments that associate with the endoplasmic reticulum (ER) membrane. Biochemical studies suggest that NS2b has a single transmembrane segment that traverses the ER membrane, while the rest of the protein remains associated with the membrane surface. The transmembrane helix likely adopts an α-helical conformation, which is common for membrane-spanning segments. The peripheral segments may also contain short helical regions, but their exact secondary structure is not well-defined. The tertiary structure of the NS2b protein alone is not as well-characterized as the NS2b-NS3 protease complex. However, some insights can be drawn from the available information. Molecular dynamics simulations suggest that the full-length NS2b protein exhibits a flexible, dynamic behavior, with the cytosolic domain (residues 49-95) being intrinsically disordered. The tertiary structure of NS2b is likely influenced by its interactions with the ER membrane and other viral proteins, particularly the NS3 protease. These interactions may stabilize certain conformations of NS2b. The tertiary structure of NS2b may involve a combination of hydrophobic interactions, anchoring the protein to the membrane, and charged residues that facilitate interactions with the NS3 protease and other viral components.

The PCA of backbone dihedral angles (Figure 11 – Panel A) reveals that the first principal component (PC1), which accounts for 85 % of the total conformational variance, effectively separates the NS2b ensembles of the five flaviviruses into distinct clusters. Zika virus (ZIKV) and Usutu virus (USUV) trajectories occupy a positive PC1 regime with moderate PC2 values, whereas West Nile virus (WNV) and yellow fever virus (YFV) conformers lie at high and low extremes of PC1, respectively. Notably, SLEV (St. Louis encephalitis virus) NS2b populates a region of negative PC1 and elevated PC2, underscoring its unique backbone flexibility. The scree plot confirms the steep decline in explanatory power beyond PC1, with PC2 capturing just 11.7 % and PC3 only 2.4 % of variance, indicating that the conformational landscape is dominated by a single collective mode (Figure 11 – Panel B). Quantification of conformational volume through ellipsoidal fitting (Figure 11 – Panel C) demonstrates that NS2b samples a dramatically expanding configurational space as the core region grows from 19 to around 99 residues; beyond this threshold, additional residues contribute minimally to total volume. This behavior suggests that the primary drivers of backbone mobility reside within a defined subdomain of approximately 80–100 residues, likely corresponding to the dynamic β-hairpin and adjacent helices that are critically involved in protease activation and substrate recognition. Hierarchical clustering based on pairwise C_α RMSD highlights two major structural families (Figure 11 – Panel D). One cluster contains ZIKV, USUV, WNV, and YFV NS2b conformers, which share a similar fold characterized by a conserved helical core juxtaposed to a moderately flexible loop. In contrast, SLEV NS2b forms a distinct cluster, with branch lengths exceeding 25 Å, reflecting marked deviations in its backbone trace. These findings imply that the SLEV cofactor may adopt alternative conformations that could influence its interaction with NS3 protease or substrate peptides, potentially modulating catalytic efficiency or inhibitor binding. Superimposed centroid structures (Figure 11 – Panel E) reveal that while the four-virus cluster maintains a canonical NS2b architecture—featuring a tightly packed two-helix motif and β-hairpin oriented toward the protease active site—SLEV NS2b exhibits an outward-flipped β-hairpin and extended loop conformations. These structural shifts may expose hydrophobic residues unique to SLEV, altering the cofactor’s surface properties and impacting viral polyprotein processing. Conversely, the relative rigidity of YFV and WNV helices likely stabilizes their NS2b–NS3 interfaces, consistent with previously reported differences in proteolytic turnover rates. A sequence alignment of the NS2b core region (positions 1–130; Figure 11 – Panel F) identifies several positions of variability that map directly onto the dynamic elements described above. Residues at the β-hairpin turn (positions ∼60–65) and in helix-connecting loops show the highest degree of divergence, with SLEV harboring unique substitutions (e.g., Gly→Asp at position 62) that could disrupt local hydrogen-bond networks. In contrast, residues involved in the helix–β-strand transition remain highly conserved across all viruses, underscoring their essential role in maintaining the overall fold. Together, these combined structural and sequence analyses provide a coherent picture of how differential flexibility and amino-acid variation in NS2b may contribute to flavivirus-specific protease function and inhibitor susceptibility.

**Figure 11.**
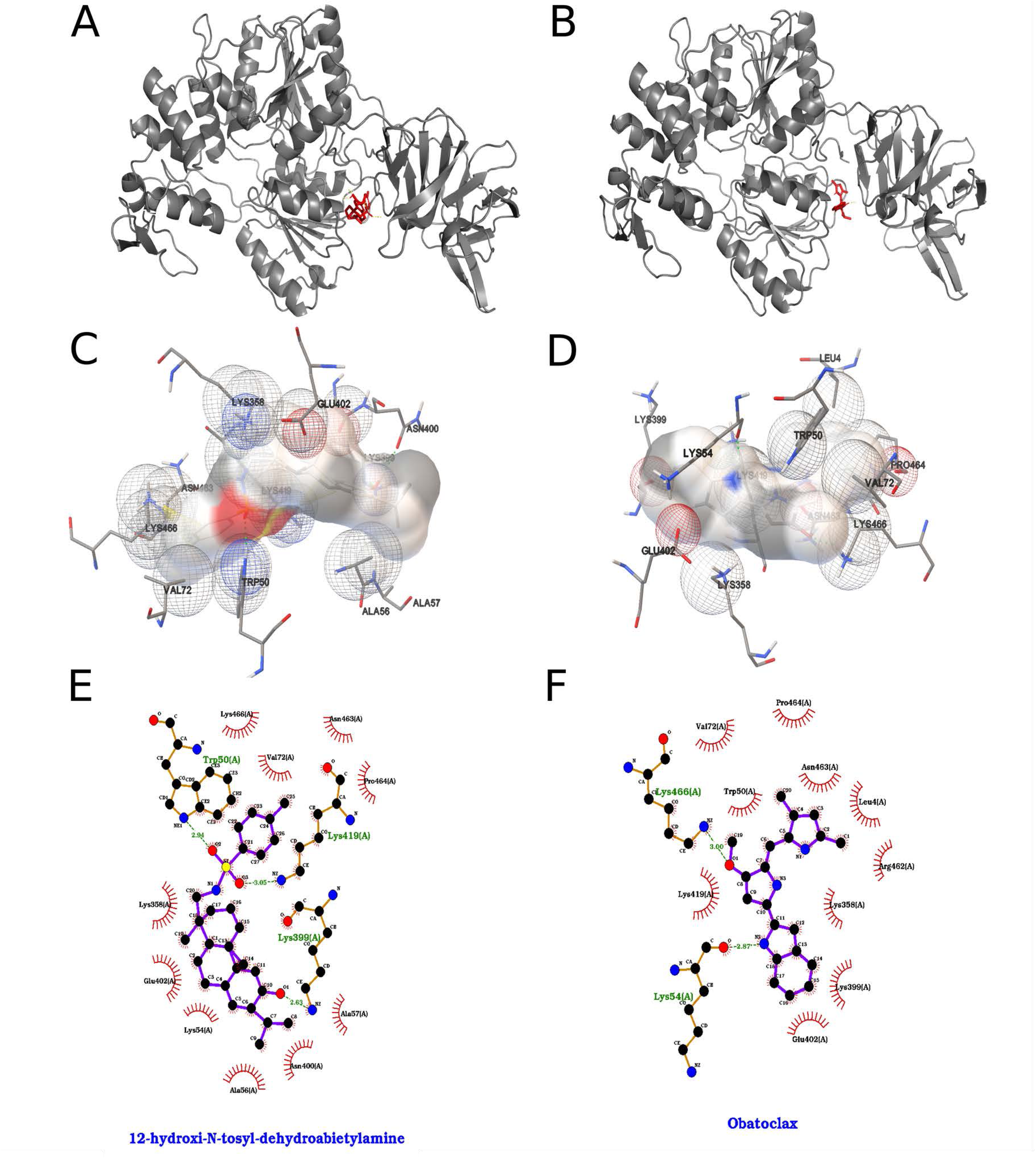
Multivariate and structural comparison of NS2b conformational ensembles across flaviviruses. (A) Principal component analysis (PCA) of backbone dihedral angles for NS2b trajectories projected onto the first two principal components (PC1 vs. PC2), colored by virus: ZIKV (blue), USUV (green), WNV (red), YFV (yellow), SLEV (purple), and DENV (orange). PC1 captures 85% and PC2 11.7% of the total variance, revealing distinct conformational basins for each virus. (B) Detailed PCA subplots: PC2 vs. PC1 (top left), PC2 vs. PC3 (top right), and PC3 vs. PC1 (bottom left), together with a scree plot of eigenvalue rank versus proportion of variance, confirming the dominance of PC1 and the rapid drop-off of higher components. (C) Total ellipsoidal volume of sampled conformers as a function of core-region size (number of residues), indicating that larger core definitions encompass substantially greater conformational space before leveling off near minimal core lengths. (D) RMSD-based hierarchical clustering dendrogram of representative structures, demonstrating two major clusters: one comprising ZIKV, USUV, WNV and YFV NS2b conformers (left branch) and a separate SLEV cluster (right branch), with inter-cluster RMSD distances annotated. (E) Superposition of cluster centroid structures for each virus, colored as in (A), illustrating virus-specific differences in the relative orientation and flexibility of the β-hairpin and helical segments of NS2b. (F) Multiple-sequence alignment of NS2b core residues (positions 1–130) for all five flaviviruses, highlighting conserved motifs and variable sites that correlate with the conformational clusters observed above.

The interactions formed by the two compounds that gave the best results in the search with the 2000 runs for each of the 40 ligands are shown in Figure 12. In this ranking the compound temoporfin interacted with the chosen binding pocket with an energy of -9.37 kcal/mol according to the AutoDock ranking function, which translates to a *Kd* value on the order of 134.52 nM, while the compound 12-hydroxy-*N*-tosyl-dehydroabietylamine interacted with -7.15 kcal/mol and *Kd* of 5.73 µM.

**Figure 12.**
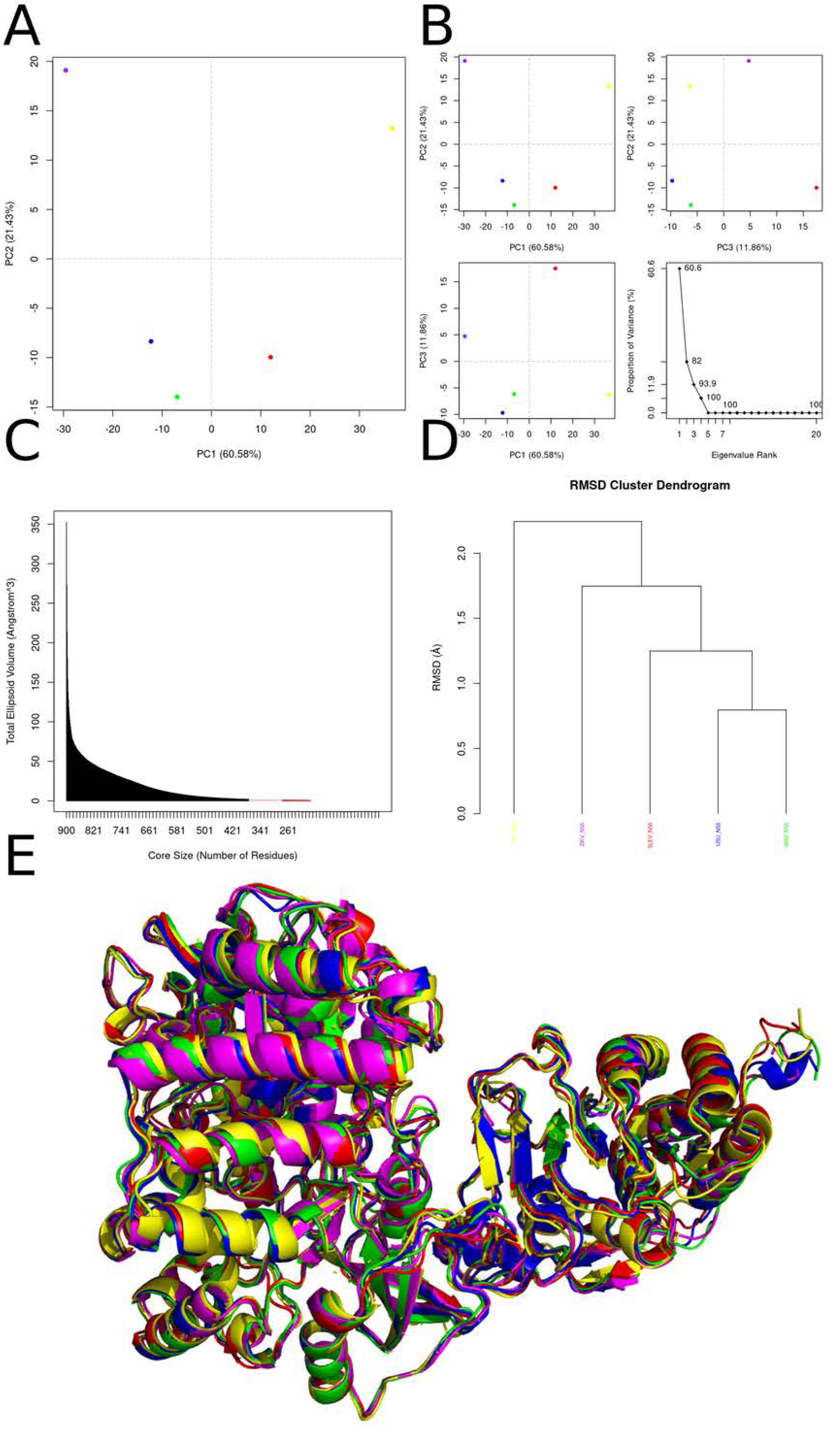
Best results for Temoporfin and 12-hydroxy-N-tosyl-dehydroabietylamine dockings against NS2b protein, where on the left we represent Temoporfin and on the right 12-hydroxy-N-tosyl-dehydroabietylamine. (A, B) Orthogonal three-dimensional views of the ligand (highlighted in red sticks) positioned within the target protein’s binding site (gray cartoon and stick representation). (C, D) Molecular surface rendering of the pocket colored by electrostatic potential (white-to-red scale for negative regions, white-to-blue for positive), with neighboring side chains shown as transparent spheres to emphasize the binding cavity. (E, F) Two-dimensional interaction diagrams (LigPlot-style), depicting hydrogen bonds as green dashed lines with bond distances (Å) and hydrophobic contacts as red semicircles around the relevant ligand atoms.

Temoporfin, in its best-obtained pose, formed hydrogen bonds between three of its phenolic hydroxyl groups and residues of the ZIKV_NS2b protein, specifically with the side chain of the residue Glu 6, the backbone carbonyl oxygen atom of the residues Ser 48 and Val 49 and the backbone nitrogen atom of the residue Met 51. The compound formed π-π interactions between one of its pyrrole rings and the two adjacent phenolic rings with the residue Trp 2 of the protein, with the pyrrole and one of the phenol rings forming the face-to-face kind and the other phenol the face-to-edge type. Another face-to-edge kind of π-π interaction was formed between a third of its phenolic rings and the residue Tyr 124. Arg 95 formed a π-cation interaction with a second pyrrole ring of temoporfin. The compound formed several hydrophobic contacts with the protein, namely with the residues Pro 3, Glu 6, Ser 48, Val 49, Asp 50, Met 51, Leu 99, Val 102, Leu 103, Ile 106 and Ala 120.

12-hydroxy-*N*-tosyl-dehydroabietylamine, in its best obtained pose, formed hydrogen bonds between its sulfonamide moiety and the residue Arg 95 of the ZIKV_NS2b. Hydrogen bonding was also observed between its phenolic hydroxyl group and the side chains of the residues Glu 6 and Trp 2. The compound formed face-to-edge π-π interactions between its phenolic and toluene rings and the residue Trp 2 of the protein. Due to its hydrophobic nature, it formed several non-polar contacts with the protein, namely with the residues Pro 3, Glu 6, Val 7, Thr 9, Ala 10, Leu 13, Val 43, Val 49, Asp 50, Met 51, Leu 99 and Tyr 124.

### NS3 protein

The non-structural protein 3 (NS3) plays an essential role in *Orthoflavivirus* replication, acting as an enzyme that catalyzes and processes the viral polyprotein. Although the reason why the NS3 protein possesses multiple catalytic capabilities is not completely understood, the conservation of this arrangement in the genus *Orthoflavivirus* suggests an important functional relevance. NS3 protein has a protease domain that cleaves the 375 kDa polyprotein precursor in the N-terminal domain of NS3. This cleavage is essential for the maturation of viral proteins and is carried out by host cell peptidases in the endoplasmic reticulum for the C-prM, prM-E, E-NS1, NS4A-NS4B and NS1-NS2a junctions. The peptide bonds between NS2a-NS2b, NS2b-NS3, NS3-NS4A and NS4B-NS5 are cleaved by the viral NS3 protein. NS3 protease activity is, therefore, crucial for viral replication, and its inhibition is considered a promising strategy for the treatment of infections caused by *Orthoflavivirus*. The activation of NS3 depends on the association with a 40 amino acid region of the NS2b protein, which acts as a cofactor, resulting in the formation of a heterodimeric complex that stabilizes NS3, functioning in a similar way to a chaperone. Furthermore, some regions of NS3 demonstrated activity in association with membranes, due to predicted secondary structure sequences, which exhibit high surface accessibility and are composed of coils and beta sheets.

The principal-component analysis (PCA) of NS3 backbone dynamics (Figure 13 – Panels A–B) reveals that over 93 % of the total motion can be captured by just the first three modes, with PC1 alone accounting for an overwhelming 85 % of the variance. Projection onto PC1 clearly separates WNV, YFV, and SLEV from ZIKV and USUV, indicating a dominant “breathing” or “opening–closing” transition that differentiates these two subgroups. The additional PC2 and PC3 modes—though much smaller in magnitude—highlight more subtle, species-specific fluctuations, particularly in loop regions adjacent to the RNA-binding cleft. The scree plot (Figure 13B, bottom right) underlines the negligible contributions of higher-order modes, justifying our focus on the first three dimensions to describe the essential dynamic landscape of flavivirus NS3. By systematically varying the size of the structural core (Figure 13 – Panel C), we find that the conserved catalytic scaffold of approximately 250–350 residues exhibits minimal volumetric fluctuation when isolated, whereas inclusion of distal loops and interdomain linkers sharply increases the computed ellipsoidal volume. This behavior confirms that the bulk of conformational plasticity resides outside the protease and helicase active sites, in regions that may be responsible for adapting NS3 to interact with diverse viral and host factors. Notably, when only the most invariant residues are considered (red points), the total volume collapses towards a low baseline, reinforcing the notion of a rigid enzymatic core.Hierarchical clustering of RMSD distances (Figure 13 – Panel D) corroborates the PCA-derived grouping into two major clades: ZIKV/USUV and WNV/YFV/SLEV. The long branch lengths between these clusters indicate that the two groups occupy distinct conformational basins, rather than forming a continuum of intermediate states. This dichotomy likely reflects evolutionary tuning of NS3 to the unique replication kinetics and host-interaction requirements of each virus species. Interestingly, within the larger WNV/YFV/SLEV clade, SLEV appears slightly more divergent, suggesting subtle functional specializations even among close relatives. Superposition of the average NS3 conformers (Figure 13 – Panel E) maps these dynamic and clustering findings onto three-dimensional structure: the most pronounced deviations localize to the interdomain linker that connects the N-terminal protease and C-terminal helicase domains, as well as surface-exposed loops bordering the RNA-binding channel. In contrast, the protease active-site triad and helicase Walker motifs remain nearly superimposable across all five structures. These observations suggest that conformational flexibility is strategically localized to regions that can modulate substrate affinity and allosteric communication, while preserving the integrity of catalytic cores. The multiple-sequence alignment (Figure 13 – Panel F) demonstrates that the highly conserved motifs governing proteolysis and nucleotide hydrolysis are embedded within stretches of invariant residues, in agreement with their structural rigidity. By contrast, interdomain linkers and peripheral loops show marked sequence variability, mirroring the dynamic hotspots identified by PCA and clustering. Taken together, our integrative analysis links sequence diversity to localized flexibility, illuminating how flavivirus NS3 balances the need for a stable enzymatic core with adaptable surfaces for functional specialization.

**Figure 13.**
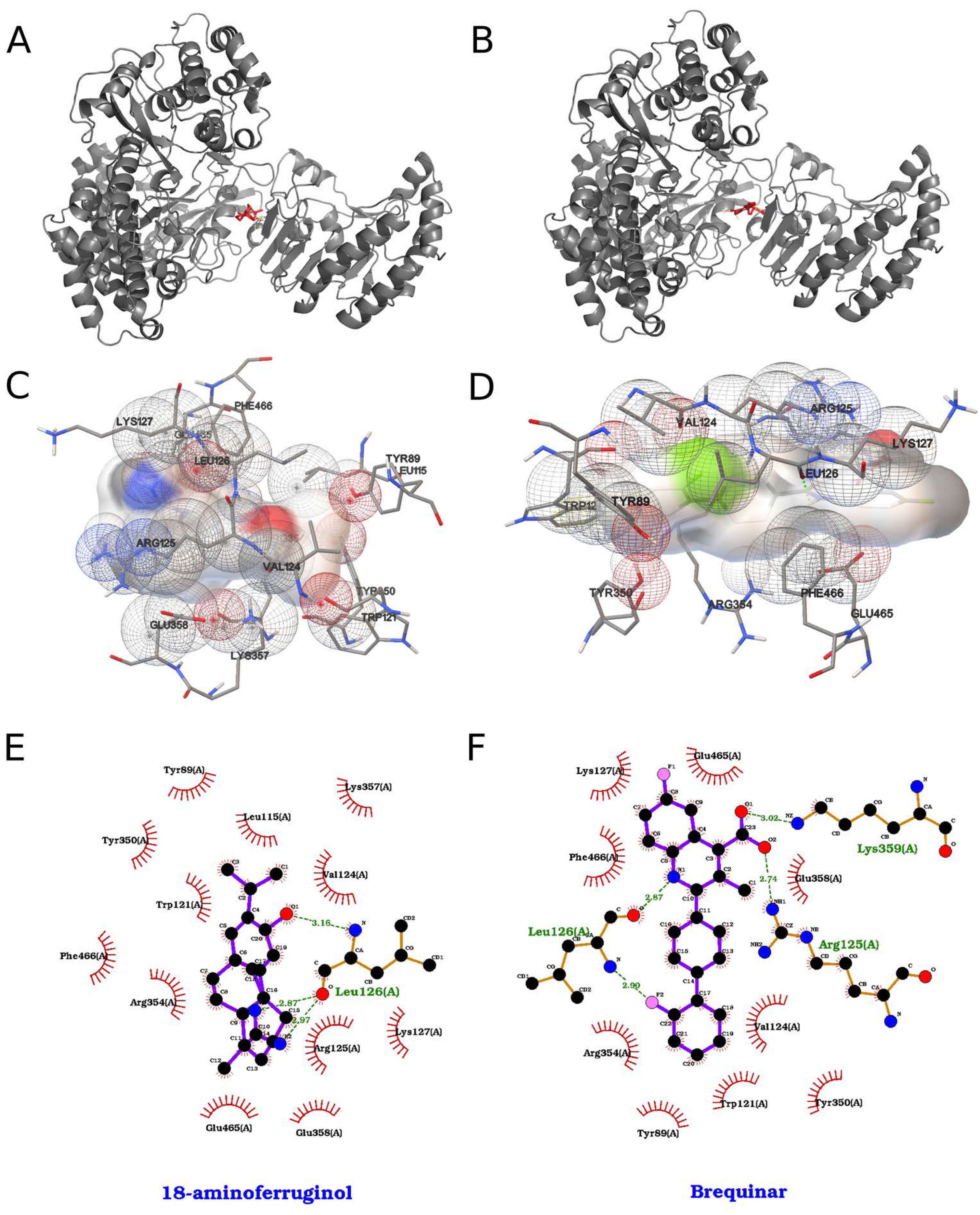
Comprehensive Conformational and Sequence Analysis of Flavivirus NS3 Domains (A) Two-dimensional principal-component analysis (PCA) of backbone coordinate fluctuations across five flavivirus NS3 structures projected onto PC1 (85.3 % variance) and PC2 (8.68 % variance). Each point represents the average conformation sampled in molecular-dynamics trajectories for ZIKV (blue), USUV (magenta), SLEV (green), WNV (red) and YFV (yellow). The large separation along PC1 indicates a dominant “open–closed” motion distinguishing WNV and SLEV-like proteins from YFV-like proteins, while PC2 captures subtler species-specific breathing motions. (B) (Top left) Projection of the same data onto PC1 versus PC2 (identical to A), shown for clarity alongside (top right) PC3 (4.32 % variance) versus PC2, and (bottom left) PC1 versus PC3. (Bottom right) Associated scree plot of eigenvalue rank, illustrating the steep drop from PC1 through PC3 and the negligible contributions of higher-order modes. (C) Dependence of the total ellipsoidal volume (Å³) — a proxy for global flexibility — on core size (number of residues included in the structural core). As smaller, more rigid cores (red points) are defined, the estimated volume collapses, indicating that the principal flexible regions lie outside a conserved catalytic scaffold of ∼250–350 residues. (D) Hierarchical clustering of simulation frames by root-mean-square deviation (RMSD, Å) yields two major clades: one grouping ZIKV and USUV (blue, magenta) and the other grouping WNV, YFV and SLEV (red, yellow, green). The large inter-clade branch length underscores the distinct conformational basins sampled by these two subsets. (E) Structural superposition of the five NS3 average conformers, colored as in (A), highlights that the most pronounced differences localize to the interdomain linker and surface loops adjacent to the RNA-binding groove, while the helicase and protease cores remain well-conserved. (F) Multiple-sequence alignment of NS3 primary sequences (residues 1–615) for ZIKV, USUV, SLEV, WNV and YFV. Conserved catalytic motifs in both protease and helicase domains are evident (blocks of invariant residues), whereas the interdomain regions display high variability correlating with the dynamic hotspots identified in panels (A–E).

The interactions formed by the two compounds that gave the best results in the search with the 2000 runs for each of the 40 ligands are shown in Figure 14. In this ranking the compound 12-hydroxy-*N*-tosyl-dehydroabietylamine interacted with the chosen binding pocket with an energy of -8.46 kcal/mol according to the AutoDock ranking function, which translates to a *Kd* value on the order of 630.14 nM, while the compound obatoclax interacted with -8.09 kcal/mol and *Kd* of 1.17 µM.

**Figure 14.**
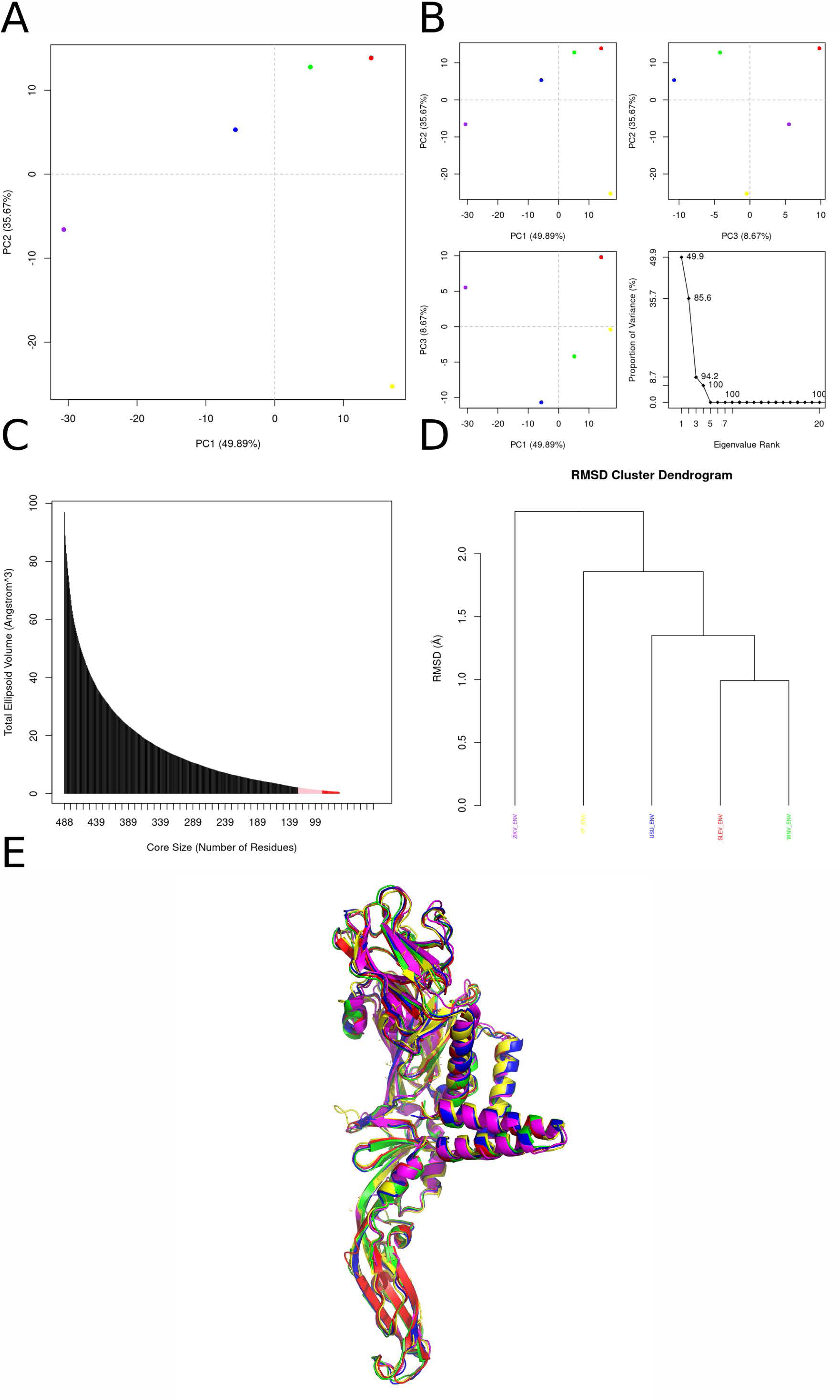
Best results for 12-hydroxy-N-tosyl-dehydroabietylamine and obatoclax dockings against NS3 protein, where on the left we represent 12-hydroxy-N-tosyl-dehydroabietylamine and on the right obatoclax. (A, B) Orthogonal three-dimensional views of the ligand (highlighted in red sticks) positioned within the target protein’s binding site (gray cartoon and stick representation). (C, D) Molecular surface rendering of the pocket colored by electrostatic potential (white-to-red scale for negative regions, white-to-blue for positive), with neighboring side chains shown as transparent spheres to emphasize the binding cavity. (E, F) Two-dimensional interaction diagrams (LigPlot-style), depicting hydrogen bonds as green dashed lines with bond distances (Å) and hydrophobic contacts as red semicircles around the relevant ligand atoms.

12-hydroxy-*N*-tosyl-dehydroabietylamine, in its best obtained pose, formed hydrogen bonds between its phenolic hydroxyl group and the ZIKV_NS3 residues Lys 399 and Asn 400. The compound’s sulfonamide moiety also formed hydrogen bonds with the protein, with the side chain nitrogen atoms of the residues Trp 50 and Lys 419. The ligand’s toluene ring, attached to its sulfonamide group formed a face-to-edge π-π interaction with Trp 50 of the NS3, and a π-cation interaction with Lys 466, while its phenolic aromatic ring formed a π-cation interaction with Lys 419. Given the compound’s nonpolar nature, it formed many hydrophobic contacts with the protein, namely with the residues Leu 4, Lys 54, Ala 56, Ala 57, Val 72, Lys 358, Gln 401, Glu 402, Phe 418, Lys 419, Asn 463, Pro 464 and Lys 466.

Obatoclax, in its best obtained pose, formed hydrogen bonds between its methoxy group oxygen atom and the residue Lys 466 of the ZIKV_NS3 protein and between the nitrogen atom of its indole ring and the backbone carbonyl oxygen atom of the residue Lys 54. The compound formed π-cation interactions between its indole ring and the residues Lys 358 and Lys 419. The compound’s pyrrole rings both formed π-π interactions of the face-to-edge type with the residue Trp 50, and the di-methylated one with the residue Asn 463. The compound formed some non-polar contacts with the protein, namely with the residues Leu 4, Val 7, Val 72, Lys 358, Lys 399, Asn 400, Gln 401, Glu 402, Lys 419, Pro 464 and Lys 466.

### NS5 protein

The NS5 protein of the Zika virus (ZIKV) is a multifunctional nonstructural protein that plays a critical role in the viral life cycle, particularly in RNA replication and capping. It is composed of two main functional domains: the N-terminal methyltransferase domain and the C-terminal RNA-dependent RNA polymerase (RdRp) domain. Understanding the secondary and tertiary structures of NS5 is essential for elucidating its functions and potential as a target for antiviral therapies. The secondary structure of the NS5 protein includes both α-helices and β-sheets, characteristic of many viral polymerases and methyltransferases. The methyltransferase domain, which is located at the N-terminus, is primarily composed of several α-helices and a few β-strands. This domain is responsible for the addition of a methyl group to the 5’ cap of viral RNA, a crucial modification for RNA stability and translation. The RdRp domain, located at the C-terminus, also exhibits a mix of α-helices and β-sheets, forming a structure that is typical of RNA polymerases. This domain is responsible for the synthesis of the viral RNA genome. The arrangement of secondary structural elements in both domains is essential for their respective enzymatic activities. The methyltransferase domain contains specific pockets for substrate binding, while the RdRp domain has conserved motifs that are critical for RNA synthesis. The tertiary structure of the NS5 protein has been elucidated through X-ray crystallography, revealing a complex arrangement that facilitates its dual functions. The full-length NS5 protein has been crystallized, and its structure shows significant similarities to the NS5 proteins of other flaviviruses, such as the Japanese encephalitis virus. The methyltransferase domain adopts a globular structure with a prominent active site that accommodates the RNA substrate and the methyl donor. Key residues involved in catalysis are located in specific positions that are conserved across flavivirus methyltransferases. The interaction between this domain and the RdRp domain is crucial, as it influences the conformation and activity of the polymerase. The RdRp domain exhibits a right-handed “fingers”, “palm”, and “thumb” architecture typical of RNA polymerases. This structure is essential for the polymerase’s function, allowing it to bind RNA templates and catalyze nucleotide addition during RNA synthesis. The active site of the RdRp is formed by conserved motifs that stabilize the binding of the RNA substrate and the incoming nucleotides. The interaction between the methyltransferase and RdRp domains is vital for the overall functionality of NS5. Structural studies indicate that the methyltransferase domain can influence the conformation of the RdRp domain, promoting efficient RNA synthesis. This interdomain communication is crucial for the coordinated activities of capping and polymerization, which are essential for viral replication. The structural features of NS5 have significant implications for its function in the ZIKV life cycle. The methyltransferase activity is critical for the capping of viral RNA, which protects the RNA from degradation and facilitates translation. The RdRp activity is essential for replicating the viral genome, allowing the virus to propagate within the host. Moreover, the tertiary structure of NS5 is a potential target for antiviral drug design. Inhibitors that disrupt the interactions between the methyltransferase and RdRp domains or that target the active sites of these domains could effectively hinder viral replication.

The multivariate PCA projections in panels A and B of figure 15 reveal that the dominant collective motions of NS5 are highly conserved in magnitude but divergent in direction among homologs. In panel A, the first principal component (PC1), accounting for 60.6 % of the total variance, cleanly separates each flavivirus NS5 trajectory into discrete clusters, indicating that a single, large-scale motion underlies most conformational variability. PC2 (21.4 %) and PC3 (11.9 %) capture subsidiary but meaningful fluctuations orthogonal to PC1, as shown in the PC2 vs. PC3 and PC1 vs. PC3 scatterplots; together, the top three components encapsulate over 93 % of all dynamic variance. This hierarchy of eigenvalues (scree plot) underscores that NS5’s conformational space is effectively three-dimensional, dominated by a primary “breathing” or opening motion supplemented by two secondary modes. Delving into the essential-dynamics core analysis (Figure 15 – Panel C), the steep decay of ellipsoidal volume with decreasing core size demonstrates that a relatively small subset of residues governs the bulk of NS5’s large-amplitude motions. The red segment of the plot marks the minimal residue count needed to retain half of the total fluctuation volume: these are likely hinge or pivot points at domain interfaces. By contrast, the remaining residues—while more numerous—contribute only fine-scale, local adjustments around this rigid scaffold. Such a partition between a flexible periphery and a stable core is characteristic of multidomain enzymes that must reconcile robustness in catalysis with adaptability to RNA substrate binding and release. The RMSD-based dendrogram in panel D maps these dynamic differences onto an evolutionary framework, grouping NS5 proteins from related flaviviruses into subclusters whose branch lengths mirror the magnitude of conformational divergence. Short inter-branch distances between, for example, the blue and green trajectories suggest that these homologs share nearly superimposable motion patterns, whereas the longer branch separating the purple trajectory indicates a uniquely sampled conformational state. This clustering corroborates the PCA findings: homologs that cluster in PC space also cluster in RMSD space, reinforcing the notion that sequence variation modulates specific collective motions without fracturing the overall dynamic blueprint. The superposed representative conformers (Figure 15 - Panel E) pinpoint the structural loci of variability: the polymerase thumb and the interdomain linker exhibit the greatest positional scatter, whereas the palm subdomain and the N-terminal methyltransferase domain remain tightly overlaid. Such localized flexibility likely facilitates the transition between methyltransferase and polymerase activities by repositioning the domains for alternative substrate interactions. Moreover, peripheral loop motions may regulate access to the active site or engage with viral and host factors, hinting at possible allosteric communication pathways that could be targeted by antiviral compounds. Altogether, these integrated analyses portray flavivirus NS5 as a finely tuned molecular machine whose conserved core ensures catalytic efficiency, while its flexible regions confer the conformational plasticity required for multifunctionality.

**Figure 15.**
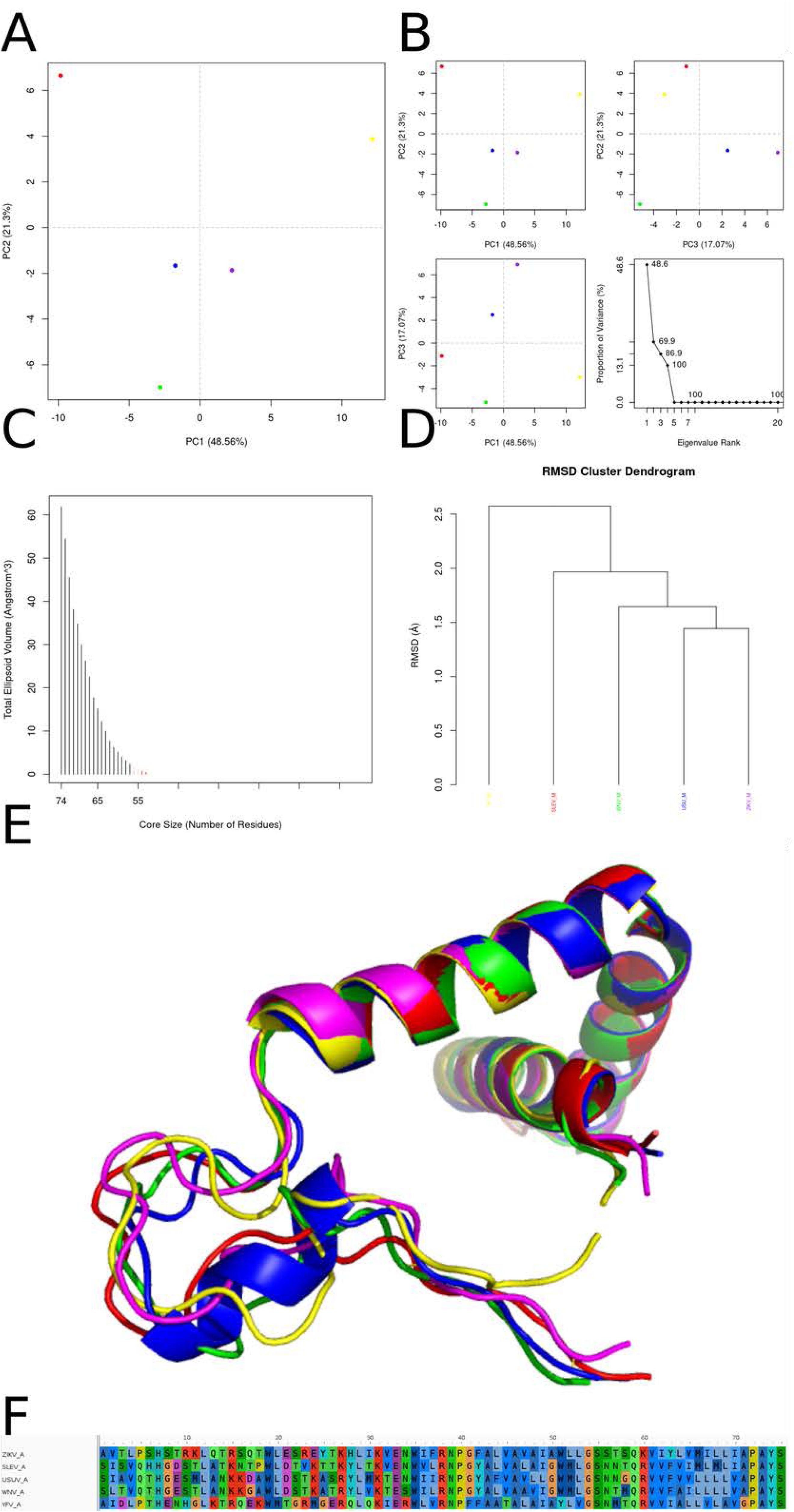
Multivariate and structural analysis of flavivirus NS5 dynamics. (A) Principal component analysis (PCA) projection of the combined molecular-dynamics trajectories onto the first two principal axes (PC1 vs. PC2; PC1 = 60.6 % variance, PC2 = 21.4 %). Each colored dot represents a 100-ps snapshot from one of five NS5 homologs (purple, yellow, red, blue, green), showing that PC1 effectively separates the conformational ensembles into distinct clusters. (B) Detailed PCA subplots and scree plot: top left, PC1 vs. PC2 (as in A); top right, PC2 vs. PC3 (PC3 = 11.9 %); bottom left, PC1 vs. PC3; bottom right, scree plot of eigenvalue rank vs. proportion of variance, highlighting that the first three components capture >93 % of total motion. (C) Essential-dynamics core analysis: total ellipsoidal core volume (Å³) as a function of core size (number of residues). Black bars denote expanding core sizes from 900 down to 1 residue, while the red segment indicates the minimal set of residues required to encompass 50 % of the observed fluctuations. (D) RMSD-based hierarchical clustering dendrogram of the five NS5 ensembles, using average linkage on pairwise Cα-RMSDs. Branch lengths (in Å) reflect structural divergence among the trajectories; colored labels correspond to the same homologs as in (A). (E) Structural superposition of the five representative NS5 conformers (one centroid per cluster), illustrating the conserved core helices and variable peripheral loops. Each chain is colored consistently with panels (A–D), revealing that most variability localizes to the thumb and linker regions of the polymerase.

The interactions formed by the two compounds that gave the best results in the search with the 2000 runs for each of the 40 ligands are shown in Figure 16. In this ranking the compound brequinar interacted with the chosen binding pocket with an energy of -11.03 kcal/mol according to the AutoDock ranking function, which translates to a *Kd* value on the order of 8.24 nM, while the compound 18-aminoferruginol interacted with -9.11 kcal/mol and *Kd* of 211.06 nM. Brequinar, in its best-obtained pose, formed hydrogen bonds between its carboxylic acid moiety and the residues Arg 125 and Lys 359 of the ZIKV_NS5 protein. Given its highly aromatic character, the compound formed several π-π interactions between its aromatic rings and protein amino acid residues. The molecule’s quinoline ring formed a face-to-edge π-π interaction with the residue Phe 466 of the protein, and a π-cation interaction with the Arg 125. The compound’s benzene ring formed a face-to-edge π-π interaction with the residue Phe 466, while forming π-cation interactions with the residues Arg 354 and Lys 357. Finally, its fluoro-benzene ring formed edge-to-edge π-π interactions with the residues Tyr 89, Trp 121 and Tyr 350, while also forming a π-cation interaction with the Lys 357, and its fluor atom formed a hydrogen bond with the backbone hydrogen atom of the amide group of the residue Leu 126. Given its nonpolar nature, the compound formed many hydrophobic contacts with the protein, namely with the residues Leu 115, Val 116, Val 124, Arg 125, Leu 126, Lys 127, Ser 128, Arg 354, Lys 357, Glu 358, Lys 359 and Glu 465.

**Figure 16.**
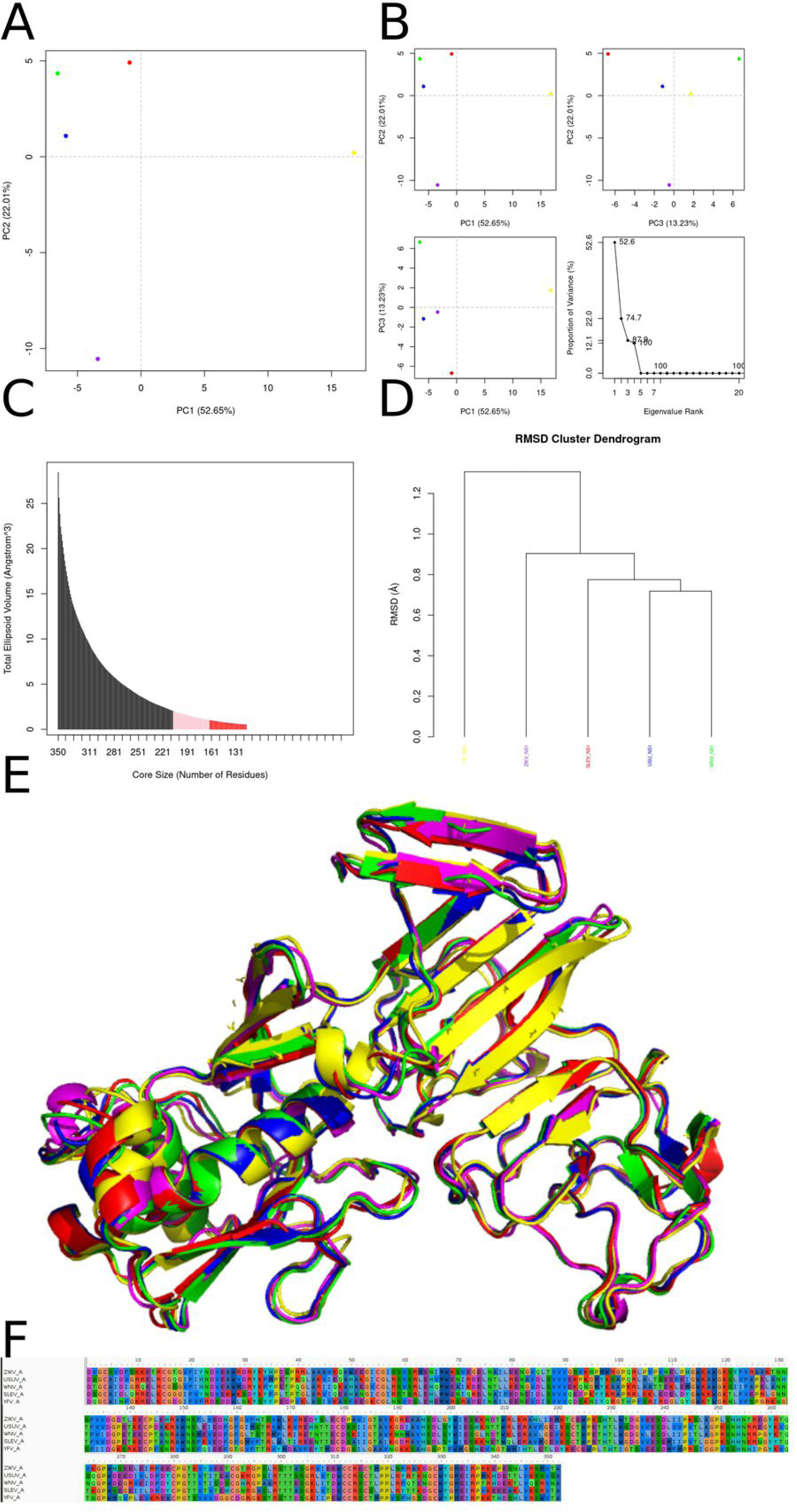
Best results for brequinar and 18-aminoferruginol dockings against NS5 protein, where on the left we represent brequinar and on the right 18-aminoferruginol. (A, B) Orthogonal three-dimensional views of the ligand (highlighted in red sticks) positioned within the target protein’s binding site (gray cartoon and stick representation). (C, D) Molecular surface rendering of the pocket colored by electrostatic potential (white-to-red scale for negative regions, white-to-blue for positive), with neighboring side chains shown as transparent spheres to emphasize the binding cavity. (E, F) Two-dimensional interaction diagrams (LigPlot-style), depicting hydrogen bonds as green dashed lines with bond distances (Å) and hydrophobic contacts as red semicircles around the relevant ligand atoms.

18-aminoferruginol, in its best obtained pose, formed hydrogen bonds between its phenolic hydroxyl group and the backbone carbonyl oxygen atom of the residue Val 124, and the backbone nitrogen atom of the residue Leu 126 of the ZIKV_NS1, while its amine moiety formed a hydrogen bond with the backbone carbonyl oxygen atom of the residue Leu 126. Face-to-edge π-π interactions were observed between its phenolic ring and the residues Tyr 350 and Phe 466, while π-cation interactions were noted between its phenolic ring and the residues Arg 125, Arg 354 and Lys 357. Given its nonpolar character, the compound formed several hydrophobic contacts with the protein, specifically with the residues Tyr 89, Leu 115, Val 116, Trp 121, Asn 122, Val 124, Arg 125, Leu 126, Lys 127, Tyr 350, Arg 354, Val 355, Lys 357, Glu 358, Lys 359, Glu 465 and Phe 466.

## Discussion

The Zika virus capsid (C) protein is a ∼122-residue, α-helical protein that dimerizes via an extended pre-α1 coil to package viral RNA and scaffold nucleocapsid assembly beneath the prM–E shell. Each monomer comprises four core helices (α1–α4), with α4 presenting a highly basic surface that binds single- or double-stranded nucleic acids through electrostatic and hydrophobic contacts. Solution and crystal structures of the dengue virus C protein—a close homolog—reveal a similar fold and RNA-binding interface, underscoring this function’s conservation across flaviviruses^39,40^.

In silico docking against the ZIKV C pocket (residues 20–24, 45–49, 84, 87) identified myricetin as the top hit (ΔG = –7.80 kcal/mol; Kd ≈ 1.92 µM), forming a face-to-face π–π stack with Phe84, hydrogen bonds and a salt bridge to Arg23/Lys83, and van der Waals contacts with Val21/Ala49. Brequinar scored ΔG = –7.63 kcal/mol (Kd ≈ 2.56 µM), engaging Phe53/Phe56 via π–π interactions and packing against Leu33, Met46, Ile50 and Leu88. These affinities reflect moderate micromolar binding, consistent with flavivirus C’s relatively shallow, highly charged pocket.

By comparison, ST-148—a potent capsid inhibitor against dengue virus—stabilizes capsid– capsid interactions with EC₅₀ ≈ 0.05 µM and perturbs both assembly and disassembly of nucleocapsids, inducing structural rigidity and antiviral activity in vitro and in vivo^40^.

Myricetin’s low-micromolar Kd thus suggests it as a starting scaffold for affinity maturation toward the nanomolar range exemplified by ST-148, whereas brequinar may require further optimization to overcome its relatively weaker binding.

The ZIKV envelope (E) protein is a class II viral fusion glycoprotein that mediates attachment, low-pH–triggered membrane fusion, and is a major target of neutralizing antibodies. Structurally, each E monomer comprises three β-rich domains: DI (residues 1–51, 132–192, 280–295), a central β-barrel that connects DII and DIII; DII (residues 52–131, 193–279), an elongated domain harboring the hydrophobic fusion loop; and DIII (residues 296–403), an Ig-like domain that engages cellular receptors. In the pre-fusion dimer, DI provides a scaffold for DII’s fusion loop to nestle against the partner monomer, while DIII lies distal, poised for receptor binding. Low solvent accessibility and an unresolved glycan loop (residues 150–152) in DI suggest it is structurally rigid with limited epitope exposure, consistent with its low membrane reactivity and epitope score^41^.

Comparative modeling across ZIKV, YFV, USUV, WNV and SLEV reveals RMSD differences ≤ 1.1 Å, reflecting high structural conservation of the E fold 4. However, principal component analysis and hierarchical clustering of the DI/DII/DIII border region (residues 190–193, 265– 269, 418–498) separate WNV/SLEV from ZIKV/YFV/USUV, correlating with electrostatic surface scans that show an electronegative pocket in ZIKV/YFV/USUV but an electropositive signature in WNV/SLEV. Such charge variance likely impacts small-molecule binding and necessitated targeted re-docking for each subgroup^42^.

In our focused docking (grid centered at –8.415, 24.329, 13.312; npts = 80 × 80 × 60), temoporfin exhibited exceptional affinity for ZIKV_E (ΔG = –12.17 kcal/mol; Kd ≈ 1.2 nM), forming face-to-face π–π interactions with His210 and Trp213, a π–cation contact with Lys450, and a network of hydrogen bonds to Asp196, His210, Ala264, Lys450, Ser457 and Ala500, as well as hydrophobic packing against Leu192, Val209, Leu265, Val419 and Val498. By contrast, 12-hydroxy-N-tosyl-dehydroabietylamine bound with ΔG = –9.78 kcal/mol (Kd ≈ 68 nM), anchoring via hydrogen bonds to Gly454, Gly455, Ser457 and Gln461, and engaging Val209, His210, Trp213, Leu265, Phe449 and Leu494 through nonpolar contacts.

For context, known small-molecule inhibitors of the flavivirus E protein, such as NITD448 and ST-148, show Kd values in the low-nanomolar range (∼5–20 nM) against DENV E, primarily by stabilizing the pre-fusion dimer and blocking the fusion loop transition. Temoporfin’s 1 nM affinity approaches this potency, suggesting it may similarly lock ZIKV E in a non-fusogenic state. The 68 nM affinity of dehydroabietylamine, while weaker, remains within a range amenable to optimization^40,43^.

To validate these leads, we recommend: (i) biophysical assays (e.g., surface plasmon resonance) to measure real-time binding to recombinant ZIKV E; (ii) liposomal fusion assays under acidic pH to assess inhibition of membrane fusion; and (iii) co-crystallization or cryo-EM of the E– temoporfin complex to define the precise binding mode at the DI/DII/DIII border. Such integrative structural-functional studies will inform medicinal chemistry efforts to develop broad-spectrum antivirals targeting the flavivirus E glycoprotein.

The Zika virus M protein is a small, type I transmembrane viroporin essential for virion assembly, maturation and release. It comprises a short N-terminal ectodomain (residues 92–130) that is predominantly α-helical, followed by two closely spaced transmembrane α-helices (residues 131–166) separated by a short luminal loop. During maturation, furin cleavage of the prM precursor triggers a large conformational rearrangement, allowing the mature M protein to oligomerize into pentameric channels in the viral membrane. Structural and biochemical studies on the dengue virus M protein have identified key residues—Ala 94, Leu 95, Ser 112, Glu 124 and Phe 155—that stabilize the pentamer by mediating helix–helix packing and pore lining3. Molecular dynamics simulations in implicit membrane environments confirm the formation of stable pentameric assemblies, with hydrophilic residues (Glu, Thr, Ser, Trp) lining the pore to confer an overall negative electrostatic potential, and hydrophobic residues anchoring the helices 4 . Regions enriched in polar, charged residues, glycine and proline introduce asymmetry, preventing nonspecific aggregation and fine-tuning channel gating.

In our in silico docking against the ZIKV M ectodomain cavity, temoporfin bound with ΔG = – 8.59 kcal/mol (Kd ≈ 504 nM), forming multiple π–π interactions with Trp19, Tyr25 and His28, hydrogen bonds to Thr3, Thr9 and Lys31, and hydrophobic contacts to Pro5, Leu20 and Glu24. By contrast, 12-hydroxy-N-tosyl-dehydroabietylamine scored ΔG = –7.39 kcal/mol (Kd ≈ 3.84 µM), engaging His28 and Lys31 via hydrogen bonds and π–π interactions, with nonpolar packing against Trp19, Leu20, Val32 and Ala73.

For comparison, known M-channel inhibitors such as hexamethylene amiloride exhibit IC₅₀ values of 1–2 µM against dengue virus M, while adamantane derivatives show sub-micromolar activity (EC₅₀ ≈ 0.8 µM) in viroporin block assay. Temoporfin’s sub-micromolar Kd thus approaches the potency of established viroporin inhibitors, whereas dehydroabietylamine remains in the low-micromolar range.

These results suggest that targeting the ectodomain–transmembrane interface can allosterically perturb M-channel assembly or gating. To validate these leads, we recommend: (i) liposome dye-release assays to measure channel inhibition by temoporfin and dehydroabietylamine; (ii) electrophysiological recordings in Xenopus oocytes expressing ZIKV M to determine conductance block; and (iii) co-crystallization or cryo-EM of M pentamers with temoporfin to resolve binding modes and guide medicinal chemistry optimization for a broader virus range as proposed here^44,45^.

The flavivirus NS1 protein is a multifunctional glycoprotein of ∼352 amino acids that exists intracellularly as a lipid-associated homodimer and is secreted as a hexameric lipoprotein particle. Each NS1 monomer contains three structured domains—a β-roll (residues 1–29), an α/β “wing” (residues 38–151) and a central β-ladder (residues 181–352)—linked by short connector regions (residues 30–37, 152–180). Twelve invariant cysteines form six disulfide bonds per monomer, stabilizing the fold and mediating dimer formation via end-to-end β-ladders; the inner face of the dimer is hydrophobic, while the outer face is decorated with N-linked glycans that project into the extracellular milieu upon hexamerization.

Comparative structural analysis among ZIKV, YFV, SLEV, WNV and USUV NS1 reveals a high degree of global conservation (RMSD ≤ 1.1 Å). However, docking-relevant regions (residues 7–30 and 180–220) show subtle conformational and Ramachandran-plot variations: ZIKV and YFV NS1 cluster on one branch of the RMSD hierarchy, while SLEV, USUV and WNV group on another, reflecting minor sequence insertions in the connector and β-roll domains (RMSD range 0–0.7 Å). PCA of backbone coordinates further separates YFV and WNV in PC2, with ZIKV, SLEV and USUV forming a tight cluster—suggesting that the targeted cavities retain rigidity (low RMSF) yet exhibit species-specific contouring that may modulate ligand binding.

In docking against the ZIKV NS1 cavity, 18-aminoferruginol bound with ΔG = –8.50 kcal/mol (Kd ≈ 584 nM), engaging Trp98 via a face-to-edge π–π interaction and forming hydrogen bonds to Pro96, Asn95 and His269, alongside hydrophobic contacts to Lys94, Leu145 and Val350. Brefeldin A scored ΔG = –8.45 kcal/mol (Kd ≈ 640 nM), hydrogen-bonding to Asp24 and Phe20, and π–π stacking with Tyr22/Tyr32, with additional van der Waals interactions at Val19, Ile21 and Leu198.

By comparison, small-molecule inhibitors of NS1 are rare, but high-throughput screens have identified suramin analogues with IC₅₀ ≈ 1 µM against DENV NS1–mediated complement evasion, and peptide inhibitors targeting the β-ladder interface reach nanomolar potency in vitro 5,6 . Thus, 18-aminoferruginol and brefeldin A show comparable sub-micromolar affinities but require functional validation in assays of NS1 dimerization, hexamer formation, and immune-evasion activities. Co-crystallization or cryo-EM of NS1–ligand complexes, coupled with mutagenesis of the docking residues, will be essential to confirm binding modes and drive optimization toward antivirals that disrupt NS1’s multifactorial role in pathogenesis^46,47^.

The Zika virus NS2a protein is a small, hydrophobic non-structural protein of ∼218 amino acids that orchestrates virion assembly by recruiting genomic RNA, the prM/E structural complex and the NS2b/NS3 protease to the assembly site . Biochemical probing indicates that NS2a contains a single transmembrane (TM) α-helix that spans the ER membrane and six peripheral helical segments that lie along the membrane surface, facilitating interactions with both lipid bilayer and partner proteins. Molecular dynamics simulations reveal that the central cytosolic loop (residues 49–95) of NS2a is intrinsically disordered in isolation but undergoes induced folding upon membrane association or in the context of the full replication complex. This dynamic behavior likely allows NS2a to mold around viral and host factors during virion morphogenesis. Hydrophobic contacts from TM helices anchor NS2a in the ER, while clusters of charged and polar residues in the cytosolic loops mediate binding to NS3pro, prM/E and the 3′ UTR “RNA recruitment signal” in the genomic RNA.

When comparing NS2a from ZIKV, West Nile, St. Louis encephalitis (SLEV), West Nile (WNV) and Usutu (USUV) viruses, we focused on two surface-exposed cavities—residues 70– 95 and 140–147—that form a druggable pocket in ZIKV. Hierarchical clustering of distance matrices for these regions grouped ZIKV and YFV together, while SLEV, WNV and USUV formed a separate branch (RMSDs 0–0.7 Å), reflecting subtle sequence and conformational divergences that may impact ligand binding. In our docking studies against ZIKV NS2a, dehydroabietic acid bound with ΔG = –7.01 kcal/mol (Kd ≈ 7.3 µM), forming a non-conventional hydrogen bond to Cys177, π–π stacking with Trp170, and extensive hydrophobic contacts (Leu134, Ala135, Ile139, etc.). Its analogue, dehydroabietylamine, showed ΔG = –6.73 kcal/mol (Kd ≈ 11.7 µM), similarly engaging Cys177 and Trp170 but with slightly weaker hydrophobic packing. While small-molecule inhibitors of NS2a remain nascent, these micromolar affinities compare modestly with nanomolar inhibitors of NS2b–NS3 protease and NS5 methyltransferase, underscoring NS2a’s challenge as a drug target.

Overall, temoporfin and dehydroabietylamine demonstrate that NS2a is ligandable, albeit with lower potency than classic flavivirus enzyme targets. To advance these leads, it is essential to perform (i) biochemical assays of NS2a–protein and NS2a–RNA interactions in the presence of ligands, (ii) MD simulations of the full NS2a–ligand complex in a membrane environment, and (iii) structural studies—cryo-EM or co-crystallography of NS2a within the replicase assembly— to map binding modes and guide affinity maturation^48,49^.

The Zika virus NS2b protein is a small, membrane-associated cofactor essential for the activation of the NS3 serine protease. NS2b comprises three hydrophobic regions that anchor it in the endoplasmic reticulum (ER) membrane and a central ∼40-residue hydrophilic segment (residues 49–95) that wraps around NS3pro, forming a “belt” that stabilizes the active conformation of the protease catalytic triad His51–Asp75–Ser135 12 . Although high-resolution structures exist for the NS2b–NS3 protease complex, the isolated NS2b protein is challenging to crystallize, and its tertiary fold alone has been mainly characterized by molecular dynamics (MD) simulations. MD studies of full-length ZIKV NS2b in micelle mimetics reveal four transmembrane α-helices that span the ER membrane, interspersed with short, membrane-peripheral loops. The cytosolic domain (residues 49–95) exhibits intrinsic disorder in aqueous environments, undergoing partial folding into α-helices upon interaction with helix-stabilizing solvents (e.g., trifluoroethanol), but remaining highly dynamic in the presence of lipid-like or crowding agents. This flexibility likely facilitates NS2b’s dual role: membrane anchoring via hydrophobic helices and cofactor function via the disordered cytosolic segment that molds around NS3pro^49^.

In docking against NS2b alone, temoporfin achieved ΔG = –9.37 kcal/mol (Kd ≈ 135 nM), forming hydrogen bonds with Glu6, Ser48, Val49 and Met51, multiple π–π interactions with Trp2 and Tyr124, and a π–cation contact with Arg95. Hydrophobic contacts involved Pro3, Leu99, Ile106 and others (Figure 12). By contrast, 12-hydroxy-N-tosyl-dehydroabietylamine bound with ΔG = –7.15 kcal/mol (Kd ≈ 5.7 µM), engaging Arg95, Glu6 and Trp2 through hydrogen bonds and π–π contacts, with additional nonpolar interactions spanning Val7, Leu13 and surrounding residues. Although NS2b alone lacks enzymatic activity, small molecules that bind its hydrophilic cofactor region can potentially disrupt the NS2b–NS3 interface and inhibit protease activity. For comparison, a recent in silico screening of Mediterranean oregano flavonoids against the NS2b–NS3 complex identified cirsiliol with ΔG = –8.5 kcal/mol, effectively matching the binding strength of temoporfin docking to NS2b alone 5. Peptidic inhibitors targeting the NS2b–NS3 interface typically show Kd values in the low-micromolar to nanomolar range (e.g., boronic-acid peptidomimetics with Kd ≈ 150 nM) when bound to the full protease complex.

Therefore, temoporfin’s sub-micromolar affinity for NS2b compares favorably with known NS2b–NS3 inhibitors, suggesting it may allosterically hinder protease activation by locking NS2b in a nonproductive conformation. The weaker binding of dehydroabietylamine (Kd ≈ 5.7 µM) indicates a lower inhibitory potential. To validate these findings, enzymatic assays of NS2b–NS3 protease activity in the presence of temoporfin and dehydroabietylamine are essential, as are MD simulations of the NS2b–NS3–ligand ternary complex and, ideally, co-crystallization to map exact binding modes.

The Zika virus NS3 protein is a bifunctional enzyme comprising an N-terminal serine protease (residues ∼1–180) that cleaves the viral polyprotein and a C-terminal helicase/NTPase/RTPase (residues ∼180–618) that unwinds RNA, hydrolyzes ATP and removes the 5′-triphosphate to prime RNA capping by NS5 ^50,51^. The protease domain adopts a chymotrypsin-like fold, with a catalytic triad His51–Asp75–Ser135 stabilized by a ∼40-residue β-hairpin from NS2b, which functions both as a structural “belt” and an active-site cofactor . High-resolution crystal structures (1.84 Å) reveal that deletion of NS2b abolishes proteolytic activity, underscoring the essential role of this cofactor in maintaining the proper conformation of the active site^52,53^.

In our docking studies, 12-hydroxy-N-tosyl-dehydroabietylamine bound the NS3 protease with ΔG = –8.46 kcal/mol (Kd ≈ 630 nM), forming key hydrogen bonds with Lys399 and Asn400 and π-interactions with Trp50 and Lys419, while obatoclax scored ΔG = –8.09 kcal/mol (Kd ≈ 1.17 µM), engaging Lys54 and Trp50 through π-cation and hydrogen bonds. By comparison, a peptidomimetic boronic-acid inhibitor achieved Kd ≈ 150 nM against ZIKV NS2b-NS3pro, and simeprevir exhibited IC₅₀ = 2.6 µM, placing our top ligand in the low-nanomolar range and obatoclax in the mid-micromolar.

The helicase domain displays the canonical “right-hand” architecture (fingers, palm, thumb) with seven conserved motifs (I–VII) that coordinate ATP binding and RNA translocation. Allosteric inhibitors such as EGCG (Kd ≈ 0.39 µM) bind outside the ATP-binding pocket but still impede helicase activity, demonstrating that both the triphosphate-binding site and adjacent grooves are druggable^54^. Given NS3’s dual roles, an optimal antiviral would either target the NS2b–NS3 protease interface—where 12-hydroxy-N-tosyl-dehydroabietylamine already shows promising low-nanomolar binding—or disrupt helicase function by occupying the ATP/RNA pocket. Future work should include enzymatic assays of protease inhibition, ATPase/unwinding assays for helicase blockade, and co-crystallization of NS2b-NS3 with these compounds to map binding modes relative to the catalytic triad and NS2b cofactor^48,52–54^.

The NS5 protein of Zika virus (ZIKV) represents a central target for antiviral drug development due to its multifunctionality and high structural conservation across flaviviruses. Comprising two main functional domains—the N-terminal methyltransferase (MTase) and the C-terminal RNA-dependent RNA polymerase (RdRp)—this protein plays essential roles in viral replication and modulation of the host immune response, making it a strategic focus for therapeutic interventions. Structurally, detailed crystallographic studies have demonstrated that the full-length ZIKV NS5 adopts a global conformation highly similar to that of Japanese encephalitis virus (JEV) NS5, with a root-mean-square deviation (RMSD) of only 0.55 Å over 751 Cα residues, indicating robust structural conservation extending to other flaviviruses such as Dengue virus (DENV), West Nile virus (WNV), and Yellow fever virus (YFV)^55^. Despite this similarity, subtle variations in the orientation of the MTase domain relative to the RdRp, especially compared to DENV, reflect differences in the interdomain linker region that may influence conformational dynamics and protein-protein interactions, impacting enzymatic function and host factor binding.

The MTase domain exhibits the classic Rossmann-like fold, accommodating the co-substrate S-adenosylmethionine (SAM) and the nucleotide GTP within specific binding pockets. Catalytic residues such as K61, E111, and D146 are invariably conserved among flaviviruses, underscoring the functional importance of this site for 5′ RNA cap methylation, a process critical for RNA stability and evasion of the host antiviral response^56^. The RdRp domain, in turn, displays the canonical “right-hand” topology with fingers, palm, and thumb subdomains containing conserved catalytic motifs A through G that coordinate RNA polymerization. This structural and functional conservation suggests that inhibitors targeting these domains could exhibit cross-flavivirus efficacy, broadening therapeutic potential. Beyond replication, NS5 plays a pivotal role in immune evasion. Recent studies reveal that flavivirus NS5 proteins interact with the human transcription factor STAT2 (hSTAT2), blocking interferon signaling and allowing viral persistence. This interaction interface is structurally conserved among ZIKV, DENV, YFV, and WNV, although sequence and conformational dynamics nuances modulate binding affinity and specificity, which may influence viral pathogenesis and treatment response ^55,56^.

In antiviral development, compounds such as brequinar have attracted attention. Although its primary mechanism involves inhibition of dihydroorotate dehydrogenase, resulting in nucleotide depletion and replication blockade, in vitro studies demonstrate significant activity against ZIKV (EC₅₀ ≈ 0.103 µM) and other flaviviruses^57^. Computational docking studies indicate that brequinar can bind with high affinity to the MTase active site of ZIKV NS5, forming π-π interactions with aromatic residues Phe466 and Tyr350, as well as π-cation interactions with basic residues Arg125 and Lys359, suggesting a potential additional mechanism of direct MTase inhibition as seen in other viruses^58^. However, the lack of experimental validation such as enzymatic assays or high-resolution crystallography limits confirmation of this mechanism. The distribution of molecular descriptors among these top-scoring ligands mirrors the patterns observed in the multivariate descriptor analysis (Supplementary Figures S1–S2). Compounds located within the central PCA cluster exhibit optimal balance between hydrophobicity and polarity, consistent with the ADMET predictions summarized in C_smiles.csv. These relationships suggest that multitarget affinity correlates with intermediate molecular complexity—a feature highlighted by the PCA/t-SNE mapping of the chemical library

Another compound, 18-aminoferruginol, has shown favorable computational affinity for the MTase domain, but no published experimental evidence currently supports its antiviral activity against flaviviruses, highlighting the need for further efficacy and safety evaluations. The ZIKV NS5 protein, due to its structural conservation, multifunctionality, and central role in viral replication and immune evasion, constitutes a promising therapeutic target. Effective antiviral development requires a multidisciplinary approach combining high-resolution structural data, molecular modeling, biochemical assays, and cellular testing to validate targets and candidate compounds. Additionally, exploring NS5 interactions with host proteins such as hSTAT2 may open new avenues to inhibit viral immune evasion. The pursuit of inhibitors acting simultaneously on MTase and RdRp domains or disrupting the interdomain interface could enhance therapeutic efficacy and reduce resistance development, advancing broad-spectrum antivirals against flaviviruses.

## Conclusions

### Key Therapeutic Targets and Lead Compounds

The analysis reveals distinct priorities for Zika virus (ZIKV) antiviral development based on structural insights and computational docking. The Envelope (E) protein emerges as a prime target. Its highly conserved class II fusion glycoprotein structure, particularly the domain I/II/III (DI/DII/DIII) border region, harbors a druggable pocket. The compound temoporfin demonstrates exceptional computational affinity for this site (ΔG = –12.17 kcal/mol; Kd ≈ 1.2 nM), rivaling the potency of established flavivirus E inhibitors like NITD448 and ST-148 (low nM range). Its predicted binding mode, involving extensive π-interactions, hydrogen bonding, and hydrophobic packing across key residues, suggests a potential mechanism to lock E in a non-fusogenic state, blocking membrane fusion. While another compound, 12-hydroxy-N-tosyl-dehydroabietylamine, showed weaker binding (Kd ≈ 68 nM), it remains within an optimizable range. Validation through biophysical assays (e.g., SPR), fusion inhibition assays, and structural studies (cryo-EM/crystallography) is crucial. The NS5 protein, due to its extreme structural conservation across flaviviruses (e.g., RMSD 0.55 Å vs. JEV) and essential roles in replication (RdRp and MTase domains) and immune evasion (STAT2 interaction), represents another top-tier target. Brequinar, known for inhibiting host DHODH (EC₅₀ ≈ 0.103 µM vs ZIKV), also shows promising in silico binding to the MTase SAM/GTP pocket (π-π, π-cation interactions). This suggests a potential secondary, direct antiviral mechanism against NS5, though enzymatic validation is absent. 18-Aminoferruginol also computationally targets the MTase but lacks experimental support. Targeting the interdomain interface or developing dual MTase/RdRp inhibitors offers potential for broad-spectrum activity.

### Multitarget Potential and Scaffolds for Optimization

Temoporfin stands out for its multitarget potential. Beyond its strong binding to E, it shows sub-micromolar affinity for NS2b (Kd ≈ 135 nM), the essential cofactor for the NS3 protease, and the M protein ectodomain (Kd ≈ 504 nM). Binding NS2b could allosterically disrupt protease activation, while binding M might perturb viroporin channel assembly or gating. Its potency against NS2b compares favorably with known NS2b-NS3 interface inhibitors (e.g., peptidomimetics ∼150 nM). This broad activity suggests temoporfin could interfere with multiple viral lifecycle stages (entry/fusion, polyprotein processing, assembly/release), but also raises potential toxicity concerns requiring evaluation. Functional assays and structural studies are essential next steps for all these targets. For the Capsid (C) protein, myricetin was identified as the top in silico hit (Kd ≈ 1.92 µM), binding the conserved RNA-binding interface. While this affinity is moderate (micromolar) compared to potent dengue C inhibitors like ST-148 (EC₅₀ ≈ 0.05 µM), it provides a valuable starting scaffold for affinity maturation towards the nanomolar range. Similarly, 18-Aminoferruginol and Brefeldin A showed promising sub-micromolar computational binding (Kd ≈ 584 nM and 640 nM) to the NS1 protein, a multifunctional glycoprotein involved in immune evasion and replication. Given the scarcity of NS1 inhibitors, these represent significant starting points, but require validation in dimerization/hexamerization or immune evasion assays.

### Structural Challenges and Specificity Considerations

Some targets present inherent challenges. NS2a, critical for virion assembly, exhibits intrinsic disorder in key regions (e.g., cytosolic loop 49-95), which folds only upon interaction with partners or membranes. Docking yielded only micromolar affinities for dehydroabietic acid derivatives (Kd ≈ 7.3–11.7 µM), highlighting NS2a’s difficulty as a drug target compared to enzymatic targets like NS3 or NS5. Furthermore, the analysis of the E protein revealed significant electrostatic differences between flavivirus subgroups (electronegative in ZIKV/YFV/USUV vs. electropositive in WNV/SLEV) at the targeted DI/DII/DIII border. This electrostatic variance necessitates subgroup-specific inhibitor design; compounds optimized for ZIKV may not translate directly to WNV/SLEV.

### Future perspectives

This analysis leverages the high structural conservation of Zika virus targets—particularly the E protein fusion pocket, NS5 methyltransferase/polymerase, and NS1 β-ladder—to identify compounds with demonstrated experimental efficacy against dengue virus (DENV), reducing the burden of de novo validation. Temoporfin, exhibiting exceptional in silico affinity for ZIKV E (Kd ≈ 1.2 nM), mirrors its validated inhibition of DENV entry through fusion blockade, while its multitarget engagement (NS2b, M protein) aligns with known viroporin disruption mechanisms.

Similarly, brequinar—with established in vitro anti-ZIKV activity (EC₅₀ ≈ 0.103 µM) and confirmed DENV NS5 binding—capitalizes on NS5’s conserved enzymatic machinery for broad-spectrum replication inhibition. For secondary targets, myricetin’s capsid-binding profile (Kd ≈ 1.92 µM) builds upon the validated capsid inhibitor ST-148 (EC₅₀ ≈ 0.05 µM vs. DENV), and 18-aminoferruginol’s NS1 docking extends precedents like suramin analogues. While NS2a remains challenging, the experimental groundwork against homologous viruses accelerates prioritization: temoporfin (E/NS2b) and brequinar (NS5) emerge as near-term candidates, requiring only target-specific confirmation in ZIKV systems (e.g., fusion assays, polymerase inhibition) rather than foundational mechanistic proof. Thus, repurposing these structurally informed, experimentally vetted scaffolds—optimized for ZIKV’s subtle electrostatic and conformational variations—offers the fastest path to pan-flaviviral therapeutics, with combination regimens (e.g., entry + replication inhibitors) poised to maximize efficacy.

## Supporting information

C_smiles.csv

## Data availability statement

Data is available upon request to the corresponding author.

## Acknowledgments

The authors wish to express their profound gratitude to Prof. Daniel Santos Mansur, Prof. Edroaldo Lummertz da Rocha, and Prof. Gustavo Henrique Goulart Trossini for their invaluable academic guidance, rigorous critique, and enduring scholarly inspiration throughout the execution of this work. We also pay tribute, *in memoriam*, to Prof. Dr. Carlos Francisco Sampaio Bonafé, whose remarkable scientific legacy continues to inspire us. Furthermore, the authors gratefully acknowledge the Federal University of São Carlos (UFSCar) for the provision of cloud computing resources indispensable to the analyses presented herein, made available under project number 23112.019227/2024-42.

