## Supplementary material for "Integrative Proteome-Wide Structural Analysis and High-Throughput Docking Identify Broad-Spectrum Antiviral Scaffolds Against Zika, Yellow Fever, West Nile, Saint Louis Encephalitis, and Usutu Viruses": Suplementary Material

### Supplementary Material — Multivariate Analysis of Molecular Descriptors

Anderson P. Soares et al.

#### Supplementary Material — Multivariate Analysis of Molecular Descriptors

This supplementary section expands the discussion presented in the main manuscript (*Integrative Proteome-Wide Structural Analysis and High-Throughput Docking Identify Broad-Spectrum Antiviral Scaffolds Against Zika, Yellow Fever, West Nile, Saint Louis Encephalitis, and Usutu Viruses*). It provides a deeper quantitative interpretation of the chemical space explored by the selected ligands, focusing on molecular complexity and physicochemical diversity across the compound library.

The analyses presented here complement the structural and docking results in the main text (Section 3: *Results and Discussion*), and readers are encouraged to consult this supplementary document for a detailed understanding of compound clustering, diversity patterns, and their implications for multitarget drug discovery.

---

##### 1. Principal Component Analysis (PCA)

A **Principal Component Analysis (PCA)** was conducted using standardized molecular descriptors extracted from the curated compound dataset. The first three components captured over **82% of the total variance**, primarily reflecting molecular size, lipophilicity (Log P), and polarity (TPSA). The 3D distribution of compounds in the PCA space (Figure S1) illustrates the overall chemical diversity, revealing distinct clusters corresponding to low-, medium-, and high-complexity scaffolds.

**Figure S1.** Three-dimensional PCA projection of the screened compounds colored according to molecular complexity. The spread along PC1 correlates mainly with molecular weight and rotatable bond count, while PC2 and PC3 capture polarity and aromatic content. Simpler scaffolds (left, red) occupy a narrow region, whereas more complex polycyclic or heteroatom-rich structures (right, blue) exhibit higher dispersion, reflecting greater topological diversity.

This dimensional reduction confirms that the selected compounds represent a chemically balanced dataset suitable for proteome-wide screening. The observed variance patterns also suggest complementary binding potential across distinct viral targets, an aspect discussed in the main manuscript.

---

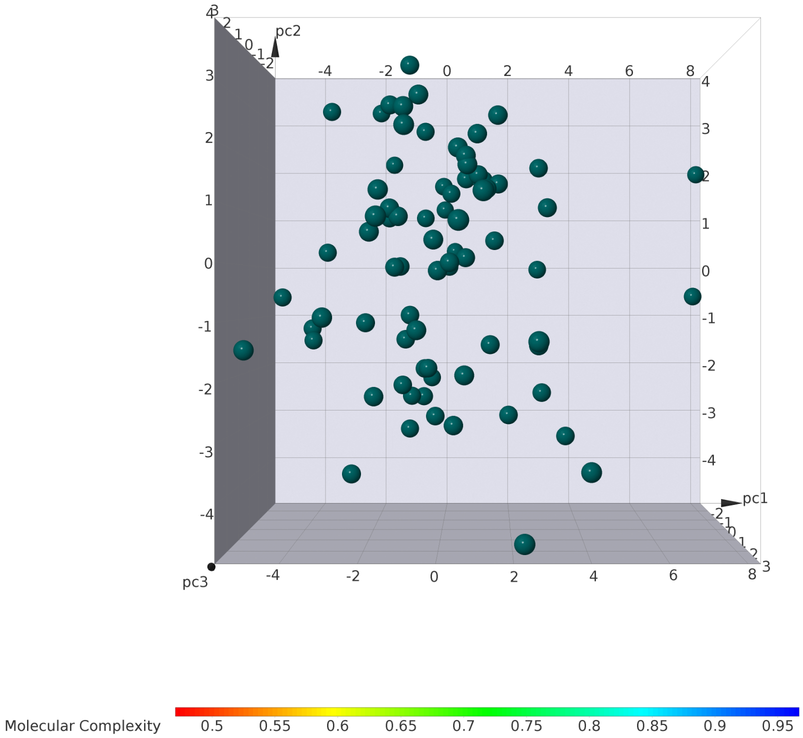

Figure 1: PCA 3D Distribution Colored by Molecular Complexity

#### 2. t-Distributed Stochastic Neighbor Embedding (t-SNE)

To better capture non-linear relationships among descriptors, a **t-SNE** embedding (perplexity = 30, learning rate = 200) was performed. The resulting 3D representation (Figure S2) shows the distribution of compounds according to **total molecular weight**, emphasizing regions of chemical space occupied by potential broad-spectrum antivirals.

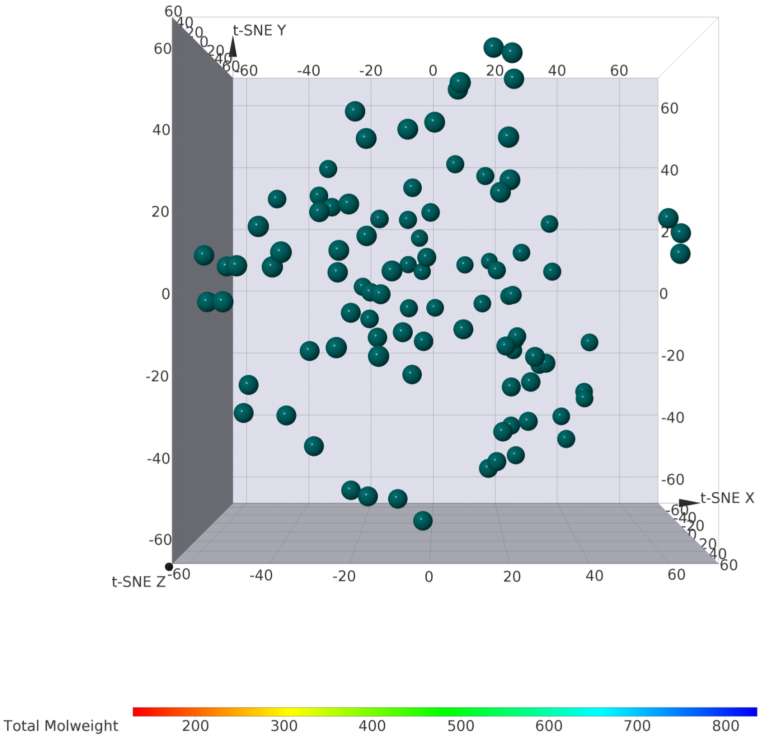

Figure 2: t-SNE 3D Distribution Colored by Total Molecular Weight

**Figure S2.** Three-dimensional t-SNE projection of the compound library, colored by total molecular weight. The relatively continuous distribution, without strong clustering, suggests that the library encompasses a wide physicochemical diversity, avoiding redundancy among analogs and covering multiple scaffold classes.

The t-SNE results complement the PCA findings by revealing subtle structural similarities not captured by linear projection. Regions of overlap between low- and medium-weight compounds indicate that physicochemical complexity does not necessarily correlate with conformational rigidity, supporting the inclusion of flexible scaffolds with balanced Lipinski profiles.

---

#### 3. Interpretation and Implications

Together, PCA and t-SNE analyses demonstrate that the compound collection used in this study occupies a **broad, non-redundant region of the antiviral chemical space**, suitable for multitarget docking and in silico optimization. The high diversity of physicochemical features reinforces the robustness of the virtual screening approach, minimizing the risk of biased target enrichment.

The main text briefly references these findings (Section 3.4, *Chemical Diversity and Ligand Selection*), where this supplementary analysis is explicitly cited as *Supporting Figure S1 and S2.* Readers seeking detailed insights into descriptor variance, clustering stability, and cross-correlations between ADME/Tox properties and structural complexity should refer to this document.

---

#### 4. Future Directions

Future analyses will integrate **UMAP embeddings** and **hierarchical clustering** of combined docking scores and ADMET metrics to identify structure–property relationships and prioritize chemical motifs with consistent multitarget affinity and favorable pharmacokinetics. These additional layers of analysis will be provided in extended supplementary materials of subsequent revisions.

---

**Corresponding Author:** Daniel F. de Lima Neto (UFSC/UFSCar)
**Supplementary Figures:** PCA 3D.png, t-SNE 3D.png
**Data Files:** C_smiles.tsv (Descriptor matrix used for multivariate analyses)
